# From climate warming to accelerated cellular ageing: an experimental heating study in a wild passerine bird

**DOI:** 10.1101/2021.12.21.473625

**Authors:** Antoine Stier, Bin-Yan Hsu, Nina Cossin-Sevrin, Natacha Garcin, Suvi Ruuskanen

## Abstract

Climate change is increasing both the average ambient temperature and the frequency and severity of heat waves. While direct mortality induced by heat waves is increasingly reported, sub-lethal effects are also likely to impact wild populations. We hypothesized that accelerated ageing could be a cost of being exposed to higher ambient temperature, especially in early-life when thermoregulatory capacities are not fully developed. We tested this hypothesis in wild great tits (*Parus major*) by experimentally increasing nest box temperature by *ca.* 2°C during postnatal growth and measuring telomere length, a biomarker of cellular ageing predictive of survival prospects in many bird species. While increasing early-life temperature had no detectable effect on growth or survival to fledging, it accelerated telomere shortening, and although non-significantly, reduced medium-term survival from 34% to 19%. Heat-induced telomere shortening was not explained by oxidative stress, but more likely by an increase in energy demand (*i.e.* higher thyroid hormones levels, increased expression of glucocorticoid receptor, increased mitochondrial density) leading to a reduction in telomere maintenance mechanisms (*i.e.* non-significant decrease in the gene expression of telomerase and protective shelterin). Our results thus suggest that climate warming can affect rate of ageing in wild birds, with potential impact on population dynamics and persistence.

## Introduction

Climate change is increasing both the average ambient temperature and the frequency and severity of heat waves (IPCC 2014). While direct mortality induced by heat waves is increasingly reported (*e.g.* [1]), sub-lethal effects are also likely to impact population dynamics and persistence [2]. Despite usually regulating their body temperature within a narrow range, endotherms can also be sensitive to (even small) changes in temperature, especially during early-life stages when their thermoregulation is not yet fully developed [3]. For instance, songbird nestlings have recently been shown to have a narrower thermoneutral zone than adults, as well as really poor cooling capacities [4]. Given that early-life experiences are known to have long-lasting effects on health, reproduction and even longevity (*e.g.* [5,6]), changes in early-life thermal environment associated with climate change are predicted to impact offspring phenotype and survival. Accordingly, some studies demonstrated associations between pre- or postnatal temperatures and survival in wild endotherm populations (*e.g.* [7–11]), and even longevity in humans [12]. Yet, the underlying physiological mechanisms remain poorly understood.

The physiological mechanisms of heat stress have been rather well characterized in laboratory endotherms: (i) Acute heat stress is known to increase the levels of key thermoregulatory and metabolic hormones, namely thyroid hormones (THs) [13,14]. (ii) Heat stress influences mitochondria, the cell powerhouse, as it reduces its efficiency to convert nutrients to ATP and increases its number [15] and (iii) increases the production of reactive oxygen species [16] contributing to oxidative stress and cellular ageing [17]. Yet, extrapolating findings from laboratory studies to wild populations in the context of climate change is challenging at best, partly because the range of temperature manipulations often exceeds temperature changes experienced in natural populations. A few studies in wild populations have experimentally increased early-life postnatal temperatures in endotherms [11,18,19], and report alterations of growth and body temperature. However, the potential mid to long-term consequences of such effects have not been characterized.

A key challenge, especially in relatively long-lived animals and wild populations, is to quantify the potential long-term deleterious effects of early-life conditions. Using the length of telomeres (*i.e.* the protective structure located at the end of chromosomes that progressively shorten with age), a key hallmark of ageing, as a molecular biomarker may help to overcome this challenge. Indeed, telomere length has been shown to predict survival (*i.e.* meta-analysis in [20]), and even lifetime reproductive success [21], and most of the telomere shortening is known to occur early in life (*e.g.* [22]). Various early-life environmental stressors, including high incubation temperature in birds (*e.g.* [22]), have been shown to accelerate telomere shortening [23]. Telomere shortening is accelerated by oxidative stress [24], mitochondrial dysfunction [25] and changes in metabolic demand [26], which are predicted to increase in response to thermal stress (see above). While thermal environment has been shown to affect telomere shortening in ectotherms (*e.g.* [27–29]), we critically lack data from endotherm species (but see [30,31]).

In this study, we comprehensively assessed the effects of an experimental increase in temperature during early postnatal development on growth, short and medium-term survival (*i.e.* to fledging and to the next autumn-winter), as well as on key physiological and ageing markers (thyroid hormones, mitochondrial density, oxidative stress, telomere length and gene expression) in a wild great tit (*Parus major*) population. The temperature elevation mimicked a scenario of *ca.* 2°C temperature increase, as forecasted by 2060 under the IPCC dangerous scenario (SSP3-7.0). It was applied during a vulnerable period of postnatal growth, *i.e.* when offspring are not anymore brooded by their mother, but still not fully capable of thermoregulation. We predicted that increasing nest temperature would lead to (i) reduced growth due to possible energetic costs of heat dissipation and/or lowered mitochondrial efficiency and/or reduced parental provisioning [32], (ii) reduced survival prospects, (iii) increased THs, (iv) increased oxidative stress, (v) increased mitochondrial density and (vi) shorter telomeres due to oxidative stress-induced shortening and/or a decrease in telomere maintenance mechanisms.

## Material & methods

### Experimental design

The experiment was conducted in 2018 in a nest box population of great tits on the island of Ruissalo (’60°26.055 N,’22°10.391 E) in Finland. Nest microclimate differs between natural nests and artificial nestboxes, and it is important to note here that we used custom-made wooden nestboxes. Great tits are mostly ectothermic until 9 days post-hatching, and then mostly endothermic from 12 days post-hatching, with a gradual transition from ectothermy to endothermy [33]. The upper critical temperature of great tit nestling increases with age (*i.e.* 6 days: 24°C; 9 days: 28°C; 12-15 days: 30°C), and the risk of lethality at high ambient temperature has been stressed previously [33]. At our field site, average temperature during the study period was mean ± SD: 15.85 ± 2.31°C and maximum daily temperature was 21.22 ± 4.43°C and was above 24.0°C for 22% of the days within the study period (data provided by the Finnish Meteorological Institute and collected by Artukainen weather station in Turku, 2 to 4 km from the study sites). Additionally, nestbox temperature is always warmer than ambient temperature, by approximately 2.5°C based on our own data (mean ± SD: 18.43 ± 2.41°C). Few breeding pairs initiate a second breeding attempt in our population, so only the first breeding attempt was used in this experimental study.

Half of the nestlings of each nest (total N =32 nests) were swapped between nests two days after hatching to account for the effects of genetic background and rearing environment. To increase nest box temperature during growth of *ca.* 2°C, one heating pad (Uniheat Shipping warmer®, USA) was installed under the ceiling in half of the nest boxes (N = 17) between d7 and d14 (hereafter, “heated” nests), *i.e.* second half of postnatal development. For the other half (N= 15), a control pad (identical to the heating pads but not producing heat) was installed (hereafter “control” nests). Heating pads were checked and replaced every second day, and control nests were also visited to standardize human disturbance between the experimental groups. The actual nest box temperature was recorded with a thermo-logger (iButton thermochron, measuring at 3min intervals, 0.0625°C accuracy) placed *ca.* 10cm above the nest cup, and daily average, minimum and maximum temperatures over the course of the heating treatment were calculated for each nest box.

Nestling body mass and tarsus length were measured prior to the treatment on d7 and post-treatment on d14 after hatching. Blood samples were collected from the brachial vein at d14 (*ca.* 70µl) using heparinized capillaries. Whole-blood samples for oxidative stress and DNA/RNA extraction were immediately snap-frozen in liquid nitrogen, and an aliquot of whole-blood was stored on ice pack until centrifugation in the laboratory at the end of the day to assess plasma thyroid hormone levels.

To study potential long-term and delayed effects of early-life heating treatment, we recaptured juvenile great tits during the following autumn-winter using mist-nets at seven feeding stations located across the field site (total of 126 hours of mist-netting). Mass and wing length of the juveniles were recorded and a blood sample (*ca.* 80µl) was collected and stored as explained above. Our method of recapture provides an estimate of post-fledging survival (*i.e.* apparent survival), but could be slightly biased by dispersal.

#### Plasma thyroid hormones and oxidative stress

Plasma thyroid hormones (T3 and T4, expressed as pg/µL) were measured from d14 nestlings with nano-LC-MS/MS following [34]. Total glutathione (tGSH), the most abundant intra-cellular antioxidant, was measured from whole-blood samples with the ThioStar® Glutathione Fluorescent Detection Kit (K005-FI, Arbor Assays, USA; technical repeatability: *R* = 0.97 (95% C.I. [0.96-0.98]). As a measure of oxidative damage, we assessed blood lipid peroxidation (malonaldehyde, MDA) using the TBARS-assay following [35] (technical repeatability: *R* = 0.92 (95% C.I. [0.88-0.94])).

### Mitochondrial density, telomere length and molecular sexing

Relative telomere length (*rTL*) and mitochondrial DNA copy number (*mtDNAcn*, an index of mitochondrial density) were quantified on DNA extracted from blood cells using real-time quantitative PCR (qPCR) assays, following [36] (see details in ESM). This technique estimates relative telomere length by determining the ratio (T/S) of telomere repeat copy number (T) to a single copy gene (SCG), and the relative mtDNAcn as the ratio between one mitochondrial gene and the same single copy gene. Birds were molecularly sexed using a qPCR approach adapted from [37,38] (see details, including on technical repeatability, in ESM).

### Gene expression analysis

We used RT-qPCR to quantify the relative expression levels of 6 genes of interest from RNA extracted from blood cells (see ESM for details on methods). We quantified the expression of genes related to (i) cellular stress response: the glucocorticoid receptor (GCR) nr3c1, two heat shock proteins of the HSP70 (*i.e.* HSPA2) and HSP90 (*i.e.* HSP90B1) families, as well as the nuclear factor erythroid 2-related factor 2 NRF2 (an oxidative-stress-induced regulator of several antioxidants and cellular protective genes); and to (ii) telomere maintenance processes: the telomeric repeat binding factor 2 TERF2 (a shelterin protein helping in protecting telomeres) and the telomerase reverse transcriptase TERT (catalytic subunit of the enzyme responsible for telomere elongation).

### Statistical analysis

For each trait of interest (see Fig. 2), we fitted a generalized linear mixed models (GLMMs, R package *lme4*) to assess the effects of heating treatment. Maximum sample size was N = 32 nests, n = 98 nestlings d14 and n = 26 juveniles, but the final sample size is often smaller and varies between physiological parameters according to sample availability and success of laboratory analyses (see specific sample size in Fig. 2). In all models, the identity of nest of origin (*i.e.* where a nestling was born) and the nest of rearing (i.e. where a nestling grew up after cross-fostering at day 2) were treated as random factors. Other random factors and fixed-effect covariates are included when relevant for the trait in question (see details in ESM tables S2-S21). Two-way interactions were initially tested, and removed from final models if non-significant (*e.g.* heating treatment * sex); yet p-value before removal are indicated in ESM tables for completeness. Main statistical outputs from type III analysis of variance with Satterthwaite’s method are reported in main text, while full model outputs with model estimates are reported in ESM tables. We report standardized effect size for each trait as Cohen’s *d* and 95% confidence interval using the *emmeans* package in *R*. For survival analyses, ln Odds Ratios were transformed to Cohen’s *d* using the formula: *d* = lnOddsRatio x (√3/∏) following [39]

## Results

The heating treatment was effective in raising average nest temperature by 1.9°C (large effect size; *F_1,30_* = 5.58, *p* = 0.025; Fig. 1) and minimum temperature by 2.3°C (large effect size; *F_1,30_* = 10.03, *p* = 0.003; Fig. 1) over the heating period, while maximum temperature was only increased by 1.3°C, which was not significant (medium effect size; *F_1,30_* = 2.31, *p* = 0.14; Fig. 1). The variance in nest temperature was significantly lower in heated nests (large effect size; *F_1,30_* = 7.02, *p* = 0.013; Fig. 1).

**Fig. 1:**
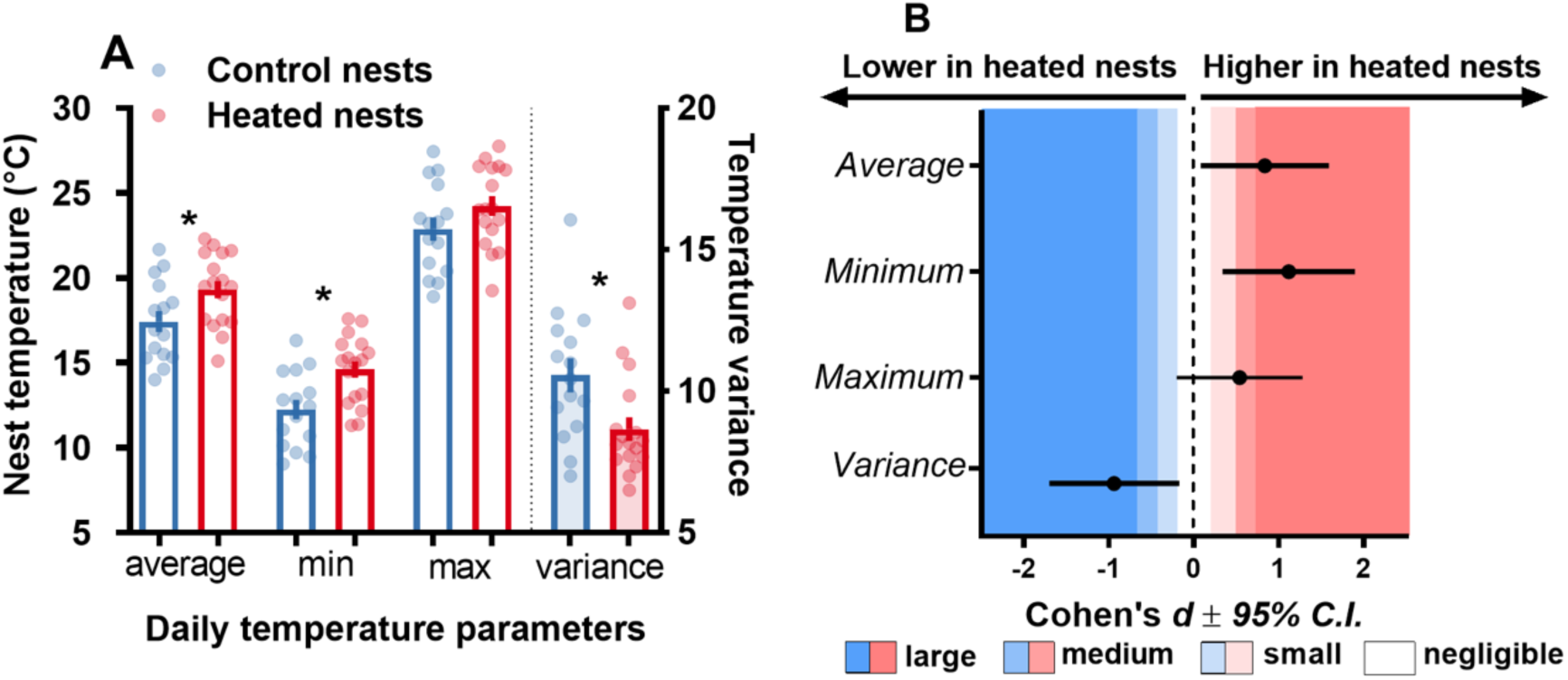
Effects of the heating treatment on nest temperature: daily average, minimum temperature, maximum and variance. (A) raw data points and mean ± SE, (B) standardized effect size and 95% confidence interval. The standard Cohen’s *d* scale for negligible <0.2, small <0.5, medium <0.8 or large >0.8 effect size is presented with a colour scale and statistical significance is indicated by an * when p < 0.05. Sample size: 15 control nests *vs.* 17 heated nests.

Nestlings from heated nests were not significantly lighter or smaller than control ones at day 14 (negligible and small effect size respectively; *p* = 0.748 and 0.969; Fig. 2, Tables S2-S3). Juveniles from heated nests were non-significantly heavier (large effect size; *p* = 0.078) but not larger (small effect size; *p* = 0.723) than control ones when recaptured the following autumn/winter (Fig. 2, Tables S4-S5).

**Fig. 2:**
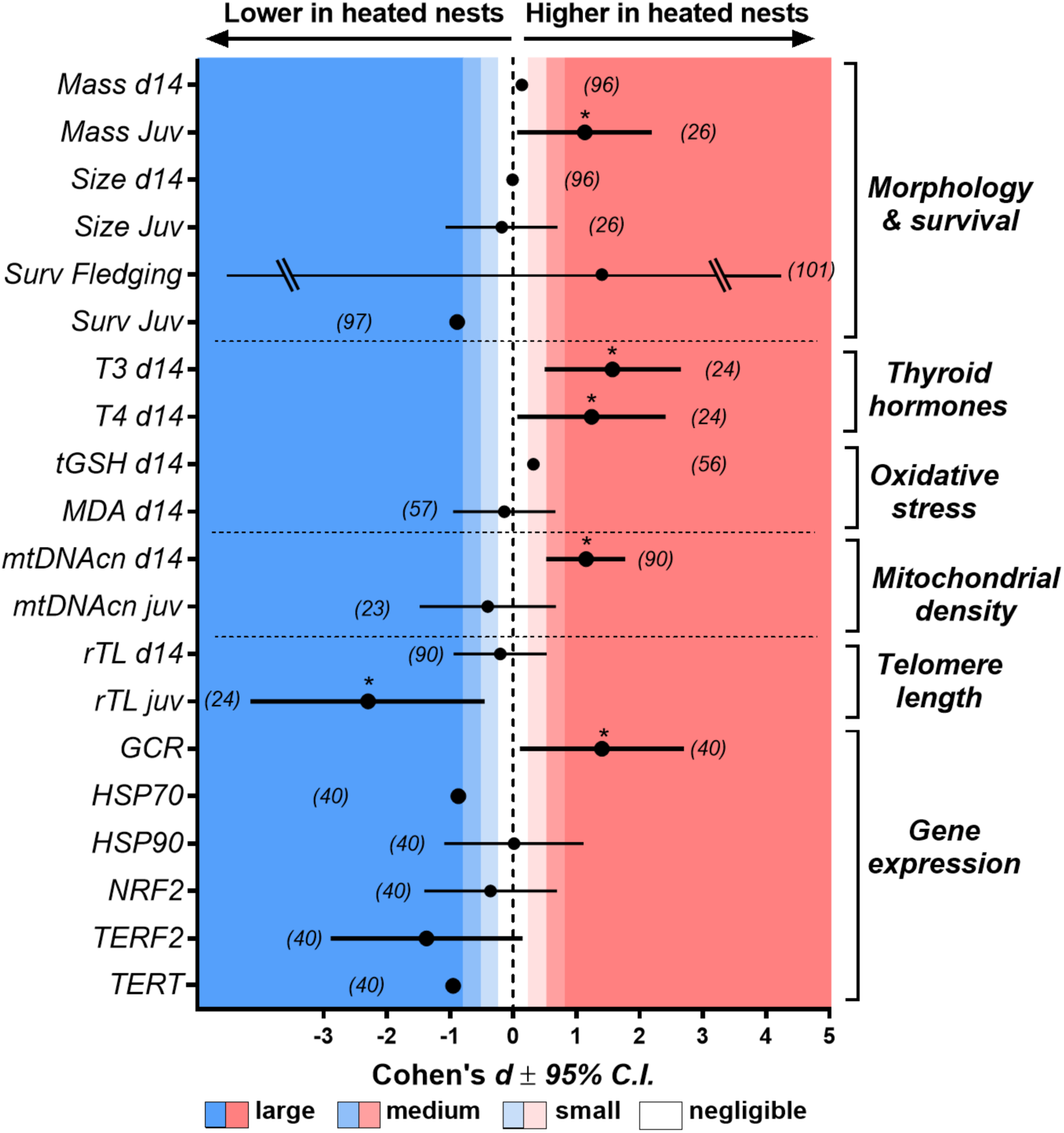
Effects of nest heating treatment on key life-history and physiological traits in wild great tits. Effects are presented as standardized effect sizes (Cohen’s *d*, or transformed to Cohen’s *d*) and 95% confidence intervals. d14: nestlings being 14 days old, Juv: juveniles (*i.e.* captured during the autumn/winter following their fledging); surv: survival; T3 and T4: plasma thyroid hormones; *MDA*: biomarker of oxidative damage to lipids measured in blood, *tGSH*: total glutathione (*i.e.* intra-cellular antioxidant molecule) content in blood cells; *mtDNAcn*: mtDNA copy number, a proxy for mitochondrial density measured in blood cells; *rTL*: relative telomere length (*i.e.* a biomarker of ageing) of blood cells; GCR: gene expression of the glucocorticoid receptor, HSP70 and HSP90: gene expression of heat shock proteins, NRF2: gene expression of an oxidative-stress-induced regulator of several antioxidants and cellular protective genes, TERF2: gene expression of a shelterin protein helping in protecting telomeres, TERT: gene expression of the catalytic subunit of the enzyme responsible for telomere elongation. Detailed information on statistics is available in ESM Tables S2-S21. Significant effects according to full statistical models are presented with * symbols, and large effects presented in bold. The standard Cohen’s *d* scale for negligible <0.2, small <0.5, medium <0.8 or large >0.8 effect size is presented with a colour scale. Sample size for each trait is presented between brackets.

Survival to fledging was very high overall and not significantly influenced by the heating treatment (control: 94% *vs.* heated: 98%; *p* = 0.82; Fig. 2, Table S6). Birds from heated nests were less likely (large effect size, control: 34% *vs.* heated: 19%;) to be recaptured as juveniles, although not significantly so: *p* = 0.22 (Fig. 2, Table S7).

Nestlings from heated nests were not significantly lighter or smaller than control ones at day 14 (negligible and small effect size respectively; *p* = 0.748 and 0.969; Fig. 2, Tables S2-S3). Juveniles from heated nests were non-significantly heavier (large effect size; *p* = 0.078) but not larger (small effect size; *p* = 0.723) than control ones when recaptured the following autumn/winter (Fig. 2, Tables S4-S5).

Survival to fledging was very high overall and not significantly influenced by the heating treatment (control: 94% *vs.* heated: 98%; *p* = 0.82; Fig. 2, Table S6). Birds from heated nests were less likely (large effect size, control: 34% *vs.* heated: 19%;) to be recaptured as juveniles, although not significantly so: *p* = 0.22 (Fig. 2, Table S7).

Nestlings from heated nests had higher circulating thyroid hormones levels than controls, significantly so for T3 (large effect size; *p* = 0.032) but not for T4 (large effect size; *p* = 0.090; Fig. 2, Tables S8-S9). Heating treatment did not significantly alter oxidative stress levels (glutathione: small effect size, *p* = 0.347; oxidative damage to lipids: negligible effect size, *p* = 0.682; Fig. 2, Tables S10-11).

Mitochondrial density in blood cells decreased sharply during postnatal development (Fig. 3A), and nestlings from heated nests had a higher mitochondrial density at the end of the heating treatment (large effect size; *p* = 0.002; Fig. 2 & 3A, Table S10), but not when recaptured as juveniles (negligible effect size; *p* = 0.554; Fig. 2 & 3A, Table S11). Heating treatment did not significantly influence nestlings’ telomere length (negligible effect size; *p* = 0.580; Fig. 2 & 3B, Table S12), but juveniles from heated nests had markedly shorter telomeres than controls (large effect size; *p* = 0.033; Fig. 2 & 3B, Table S13).

**Fig. 3:**
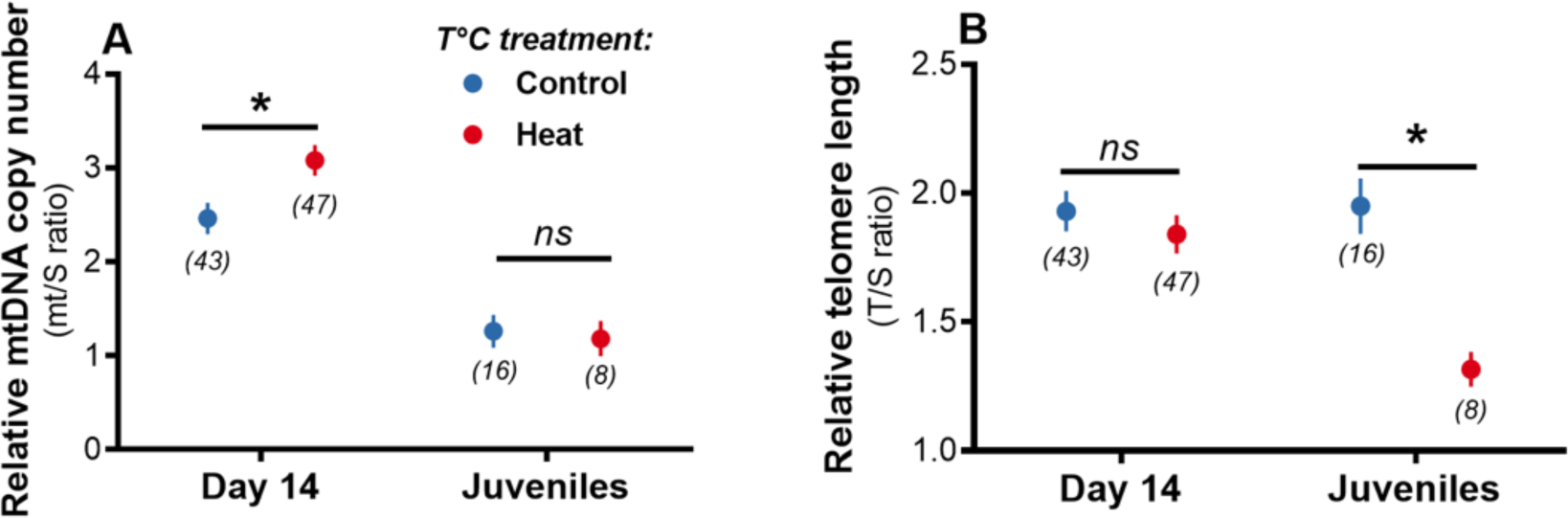
Effects of age and nest heating treatment on: (A) mtDNA copy number as a proxy of mitochondrial density and cellular energetics, and (B) relative telomere length as a biomarker of ageing. Means are plotted ± SE and full statistical models are presented in Tables S12-S15.

Gene expression was partly altered by nest heating treatment, with nestlings from heated nest showing significantly higher expression levels of the glucocorticoid receptor (GCR: large effect size; *p* = 0.045; Fig. 2, Table S16), and although non-significantly, lower expression levels of the telomere-related genes TERF2 (large effect size; *p* = 0.088; Fig. 1, Table S17) and TERT (large effect size; *p* = 0.085; Fig. 1, Table S18). There was however no clear effect on the expression of heat shock proteins (HSP70: medium effect size, *p* = 0.298; HSP90: negligible effect size, *p* = 0.985; Fig. 2, Tables S19-20) or the regulator of antioxidant protection NRF2 (small effect size, *p* = 0.504; Fig. 2, Table S21).

## Discussion

By experimentally manipulating early postnatal temperature of *ca.* 2°C (in line with predictions of climate change), we demonstrated that exposure to higher ambient temperature during early-life could affect several physiological pathways (*i.e.* thyroid hormones, mitochondrial biogenesis, glucocorticoid signalling) and ultimately lead to accelerated cellular ageing (*i.e.* shorter telomeres) through a potential deregulation of telomere maintenance processes (*i.e.* shelterin protein and telomerase). While immediate survival was not impacted by the experimental increase in nest temperature, birds from heated nests were less likely to be seen alive the following autumn-winter, although non-significantly so.

Growth rate is known to be influenced in a complex manner by weather conditions [40], and previous experimental studies increasing nest temperature have reported contrasted results (positive effect: [18]; negative effects: [11,19]). Our experimental manipulation did not affect body size or mass during postnatal growth. Apparent survival to autumn-winter was lower for birds originating from heated nests (19% *vs.* 34%, large effect size), but not significantly so. Obtaining large enough sample size to detect significant effects on survival in experimental studies on wild animals is unfortunately challenging. Yet, our results are in accordance with observational data from a 12-year monitoring study of great tit survival in Spain, showing a negative association between ambient temperature during postnatal growth and post-fledging survival [7].

While there is some evidence from laboratory acute heat stress studies that thyroid hormones could be up-regulated when facing increased ambient temperatures [13,14], we show here that even a small increase in early-life temperature can increase thyroid hormones levels. Importantly, high thyroid hormone levels have been linked to increased mortality risks in adult humans [41] and increased susceptibility to free radicals in birds [42]. Yet, we found no significant effect of nest heating on two oxidative stress biomarkers (glutathione and oxidative damage to lipids), nor on NRF2 gene expression (an oxidative-stress-induced regulator of several antioxidants). Similarly, the gene expression of two heat shock proteins (HSP70 and HSP90 families) was not clearly impacted by nest heating. This suggests that a *ca.* 2°C increase in temperature might be too low to induce oxidative stress or a heat shock response (compared to heat stress experiments in the lab that often use > +10°C; *e.g.* [43]). Yet, the gene expression of the glucocorticoid receptor nr3c1 was increased in heated nests, suggesting either a response to a stressful stimulus (*e.g.* [44]) or potentially an increase in metabolic demand [45].

Both the rise in thyroid hormones and the observed increase in mitochondrial density at the end of the heating period would support the hypothesis of an increase in metabolic demand. In line with our results on mitochondrial density, mild heat stress has been shown to increase mitochondrial biogenesis *in vitro* [15] and *in vivo* [46]. This could be a way to compensate the typical decrease in mitochondrial coupling efficiency observed at higher body temperature (*e.g.* [47]), but measuring both body temperature (*i.e.* [11] showed that nest heating induced hyperthermia) and mitochondrial coupling efficiency [48] would be needed here to test this hypothesis.

While telomeres were not immediately impacted by the heating treatment (*i.e.* no effect at the end of the heating period on day 14), we discovered some important delayed effect since juveniles from heated nests had markedly shorter telomeres than control individuals. Sample size of juveniles was relatively limited, and the lower apparent post-fledging survival of birds from heated nests could lead to bias associated with selective disappearance. Yet, this is unlikely to bias our results since we would expect individuals with shorter telomeres to disappear earlier from the population (and not the opposite), as this has previously been shown in our model species [49]. Additionally, the trend towards a decreased expression of genes related to telomere maintenance (TERF2 and TERT) observed at day 14 in nestlings from heated nests support the results observed on telomere length in juveniles. Our results are in accordance with recent findings in humans showing: 1. a negative effect of warm temperature and a positive effect of cold temperature during gestation on newborn telomere length [50]; 2. shorter telomeres in adults exposed to acute (i.e. 1-13 days) warm environmental conditions [51]. Additionally, it has recently been shown at the correlative level that warm and dry conditions were associated with shorter telomere in an Australian passerine species [30]. Here, we provide experimental evidence suggesting a causal link between environmental temperature and telomere shortening, as well as some hints on the underlying molecular mechanisms. While oxidative stress-induced telomere shortening [24] is unlikely to explain our results (see above), the ‘metabolic telomere attrition hypothesis’ [26] could be a good candidate. Indeed, this theory stipulates that increased metabolic demand mediated by increased glucocorticoid signalling (*i.e.* increased nr3c1 expression observed here) can decrease investment in telomere maintenance processes (*i.e.* decreased expression of TERF2 and TERT observed here) and be associated with mitochondrial dysfunction [52].

To conclude, our study provides the first experimental evidence that a moderate increase in ambient temperature during early-life can accelerate cellular ageing in a wild endotherm species. While the exact consequences of such increase in early-life temperature on individual fitness and population dynamics remain to be determined, previous evidence at the inter-population level has shown that shorter telomeres might precede population extinction and be used as an early warning sign [53].

## Funding

This study was financially supported by the Academy of Finland (#286278 to SR). NCS acknowledges support from the EDUFI Fellowship and Maupertuis Grant. B-Y.H work was supported by a grant from the Ella and Georg Ehrnrooth Foundation. AS was supported by a ‘Turku Collegium for Science and Medicine’ Fellowship and a Marie Sklodowska-Curie Postdoctoral Fellowship (#894963). All authors declare no conflict of interest.

## Acknowledgements

We are grateful to Jorma Nurmi, Lucas Bousseau, Thomas Rossille, Hsiao-Yin Liu and Päivi Kotitalo for their help in the field. The laboratory facilities were supported by Biocenter Finland

## Data accessibility

All data are publicly available on FigShare: 10.6084/m9.figshare.28838297

## Ethical permits

All procedures were approved by the Animal Experiment Committee of the State Provincial Office of Southern Finland (license number ESAVI/2902/2018) and by the Environmental Center of Southwestern Finland (license number VARELY549/2018) granted to SR.

## qPCR assays for telomere length, mtDNA copy number and molecular sexing

We extracted DNA from blood cells using a standard salt extraction alcohol precipitation method ([1]). Extracted DNA was diluted in elution buffer BE for DNA preservation. DNA concentration and purity (260/280 > 1.80 and 260/230 > 2.00) were checked with a ND-1000-Spectrophotometer (NanoDrop Technologies, Wilmington, USA). DNA integrity was verified in 24 samples chosen randomly using gel electrophoresis (50 ng of DNA, 0.8 % agarose gel at 100 mV for 60 min) and DNA staining with Midori Green. Each sample was then diluted to a concentration of 1.2 ng.µL^-1^ for subsequent qPCR analysis.

Relative telomere length (*rTL*) and mitochondrial DNA copy number (*mtDNAcn*, an index of mitochondrial density) were quantified using qPCR. Here, we used recombination activating gene 1RAG1 as a single copy gene (verified as single copy using a BLAST analysis on the great tit genome) and cytochrome oxidase subunit 1 (COI1) as a mitochondrial gene (verified as non-duplicated in the nuclear genome using a BLAST analysis). The qPCR reactions were performed on a 384-QuantStudio™ 12K Flex Real-Time PCR System (Thermo Fisher), in a total volume of 12µL including 6ng of DNA, primers at a final concentration of 300nM and 6μL of SensiFAST^TM^ SYBR lo-ROX (Bioline). Telomere, RAG1 and COI2 reactions were performed in triplicates on the same plates (10 plates in total); the qPCR conditions were: 3min at 95°C, followed by 35 cycles of 10 s at 95°C, 15 s at 58°C and 10s at 72°C. A DNA sample being a pool of DNA from 10 adult individuals was used as a reference sample and was included in triplicate on every plate. The efficiency of each amplicon was estimated from a standard curve of the reference sample ranging from 1.5 to 24ng. The primer sequences, as well as qPCR efficiencies and technical precision estimates (coefficient of variation and technical repeatability) are provided in Table S1. The relative telomere length and mtDNAcn of each sample were calculated as (1+*Ef*_Tel or COI2_)^ΔCq Tel or COI2^/(1+*Ef*_RAG1_)^ΔCqRAG1^; *Ef* being the amplicon efficiency, and ΔCq the difference in Cq-values between the reference sample and the focal sample.

The use of mtDNAcn as an index of mitochondrial density has been questioned in human [2], but we have previously shown good correlations between mtDNAcn and mitochondrial respiration rates in pied flycatcher [3] and great tit (Cossin-Sevrin, et al. *in revision*). Great tits have quite peculiar telomeres, characterized notably by some ultra-long telomeres that do not seem to shorten with age in adults [4]. Since qPCR only provides an estimate of overall telomere length, it could be suboptimal for this study species. Yet, relative telomere length (*i.e*. measured using qPCR) in this species has been shown to shorten during the nestling stage [5,6], to respond to environmental factors (e.g. hatching asynchrony: [5]; altitude: [6]; urbanization: [7]) and to predict adult survival [8]. Within-individual repeatability of telomere length has recently been suggested to be an important factor to evaluate the pertinence of telomere length data in a given study/species [9], and the biological repeatability in our dataset was *R* = 0.44 [0.25-0.60], which is close to the average reported by qPCR studies (*i.e. R* = 0.47), and well within the range of what has been reported for great tits [9].

Birds were molecularly sexed using a qPCR approach adapted from [10,11]. Forward and reverse sexing primers were 5ʹ- CACTACAGGGAAAACTGTAC-3ʹ (2987F) and 5ʹ- CCCCTTCAGGTTCTTTAAAA −3ʹ (3112R), respectively. qPCR reactions were performed in a total volume of 12µL including 6ng of DNA, primers at a final concentration of 800nM and 6μL of SensiFASTTM SYBR® Lo-ROX Kit (Bioline). qPCR conditions were: 3 min at 95°C, followed by 40 cycles of 45 s at 95°C, 60 s at 52°C and 60s at 72°C, then followed by a melting curve analysis (95°C 60s, 45°C 50s, increase to 95°C at 0.1°C/s, 95°C 30s). Samples were run in duplicates in a single plate and 6 adults of known sex were included as positive controls. Sex was determined by looking at the dissociation curve, with two peaks indicating the presence of a Z and W chromosome (female), and one peak indicating the presence of only the Z chromosomes (male).

## RT-qPCR assays for evaluating gene expression

We used RT-qPCR to quantify the expression levels of 6 genes of interest in d14 nestlings (Table S1). First, RNA was extracted from 10 μL of blood (immediately after the first thawing following −80°C storage) using Nucleospin RNA Plus extraction kit (Macherey-Nagel) following manufacturer instructions. Second, RNA concentration and purity were quantified using optical density. Samples not meeting quality criteria (i.e. RNA concentration > 25 ng/μl, 260/280 and 260/230 > 1.80) were excluded for further analysis. RNA integrity was checked using E-Gel 2% electrophoresis system (Invitrogen), and the ribosomal RNA 18S *vs.* 28S bands intensity, and deemed satisfactory. Samples were stored at −80°C for 2 weeks before cDNA synthesis. 600ng of RNA were used for cDNA synthesis using the SensiFAST^TM^ cDNA Synthesis kit (Bioline) following manufacturer instructions. cDNA was diluted at a final concentration of 1.2 ng/μL for qPCR analysis. No-RT control samples were prepared following the same protocol, but without reverse transcriptase enzyme.

We assessed the expression genes related to 1) cellular stress response: the glucocorticoid receptor (GCR) nr3c1, two heat shock proteins of the HSP70 (*i.e.* HSPA2) and HSP90 (*i.e.* HSP90B1) families, as well as the nuclear factor erythroid 2-related factor 2 NRF2 (an oxidative-stress-induced regulator of several antioxidants and cellular protective genes); and 2) telomere maintenance processes: the telomeric repeat binding factor 2 TERF2 (a shelterin protein helping in protecting telomeres) and the telomerase reverse transcriptase TERT (catalytic subunit of the enzyme responsible for telomere elongation). qPCR was performed in a total volume of 12μl containing 5μl of each diluted cDNA sample (i.e. 1.2ng/μl) and 7μl of reaction mix containing primers (forward and reverse) at a final concentration of 300nM and Sensifast SYBR®No-ROX Mix (Bioline). The succinate dehydrogenase complex subunit A (SDHA; [12]) and the ribosomal protein L13 (RPL13, [13]) were used as reference genes. SDHA and RPL13 were identified as the most stable reference genes using geNorm software (geNorm M < 0.7, geNorm V < 0.15) and the geometric mean of these two genes was thus used as our reference gene. Primers (Table S1) have been designed whenever possible on exon-exon junction using NCBI primer designing tool using the *Parus major* reference genome. Specificity has been checked using BLAST analysis and confirmed by a single narrow peak in melting curve analyses and the presence of a single PCR product of the expected size on agarose gel. Amplification in no-RT controls never occurred before at least 5 cycles after the lower Cq sample (> 8 Cq for all genes but TERT), and thus contamination by genomic DNA could not interfere with our results.

We ran gene-specific qPCR plates and each sample was analyzed in duplicate. Experimental groups were always balanced within each plate. We used a cDNA reference samples (*i.e.* ratio = 1) being a pool of 5 different individuals on every plate. One inter-plate standard sample was also run on every plate. qPCR assays were performed on a Mic qPCR instrument (Bio Molecular Systems) and included a two-step cycling with the following conditions: 2 minutes at 95°C; then 40 cycles of 5s at 95°C followed by 20s at 60°C (fluorescence reading) for all reactions. The expression of each gene was calculated as (1+Ef_Target_)^ΔCq(Target)^/ geometric mean [(1+Ef_SDHA_)^ΔCqSHDA^; (1+Ef_RPL13_)^ΔCqRPL13^], Ef being the amplification’s efficiency and 1′Cq being the difference between the Cq-values of the reference sample and the sample of interest.

---

**Table S1:**
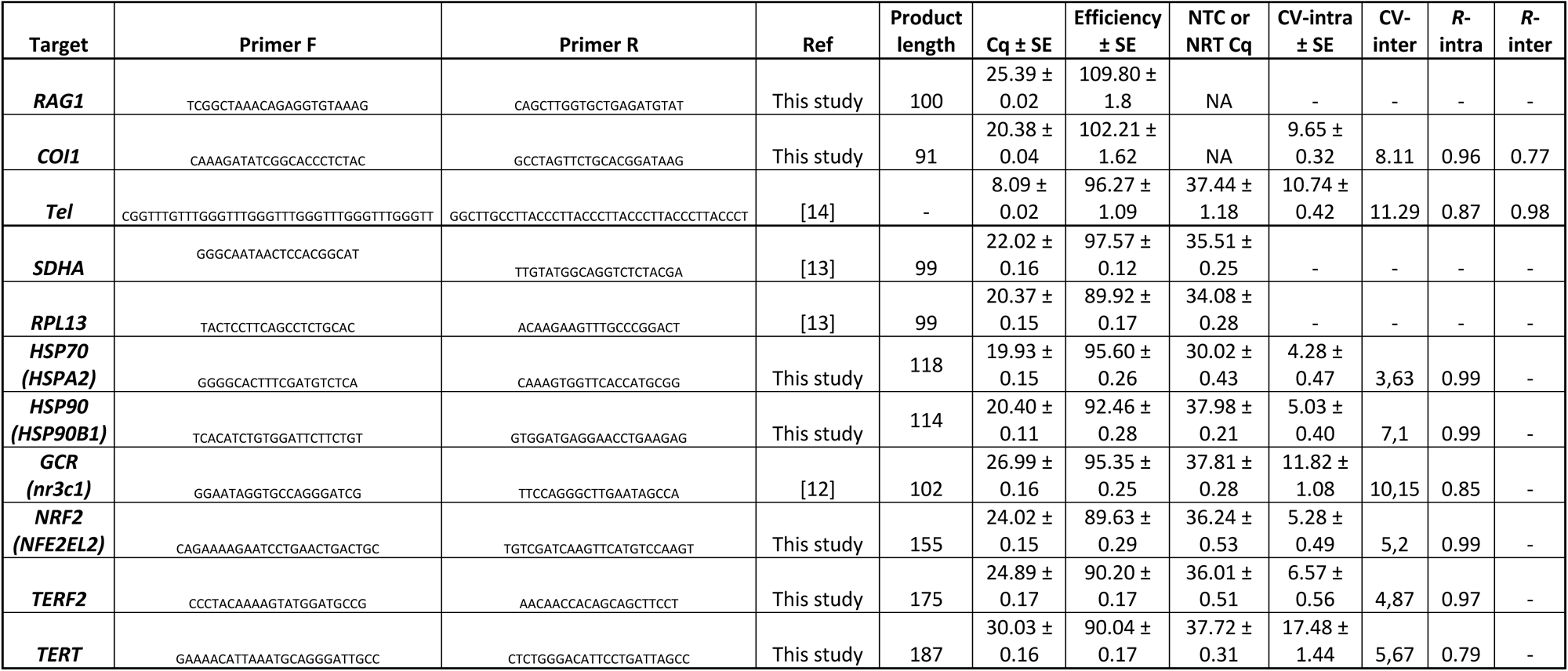
Information on qPCR primer sets and performance for relative mtDNA copy number, telomere length and gene expression assays. Cq refers to qPCR quantification cycle, efficiency has been evaluated using a standard curve for genomic DNA qPCR and using the LinReg method based on individual well efficiency for RT-qPCR for gene expression. Cq of non-template control (NTC) for genomic DNA qPCR and of non-reverse transcription control (NRT) for RT-qPCR are provided. Technical precision estimates (coefficient of variation CV and technical repeatability *R*) are provided for final ratios both at the intra-plate (based on duplicates) and the inter-plate levels.

**Table S2:**
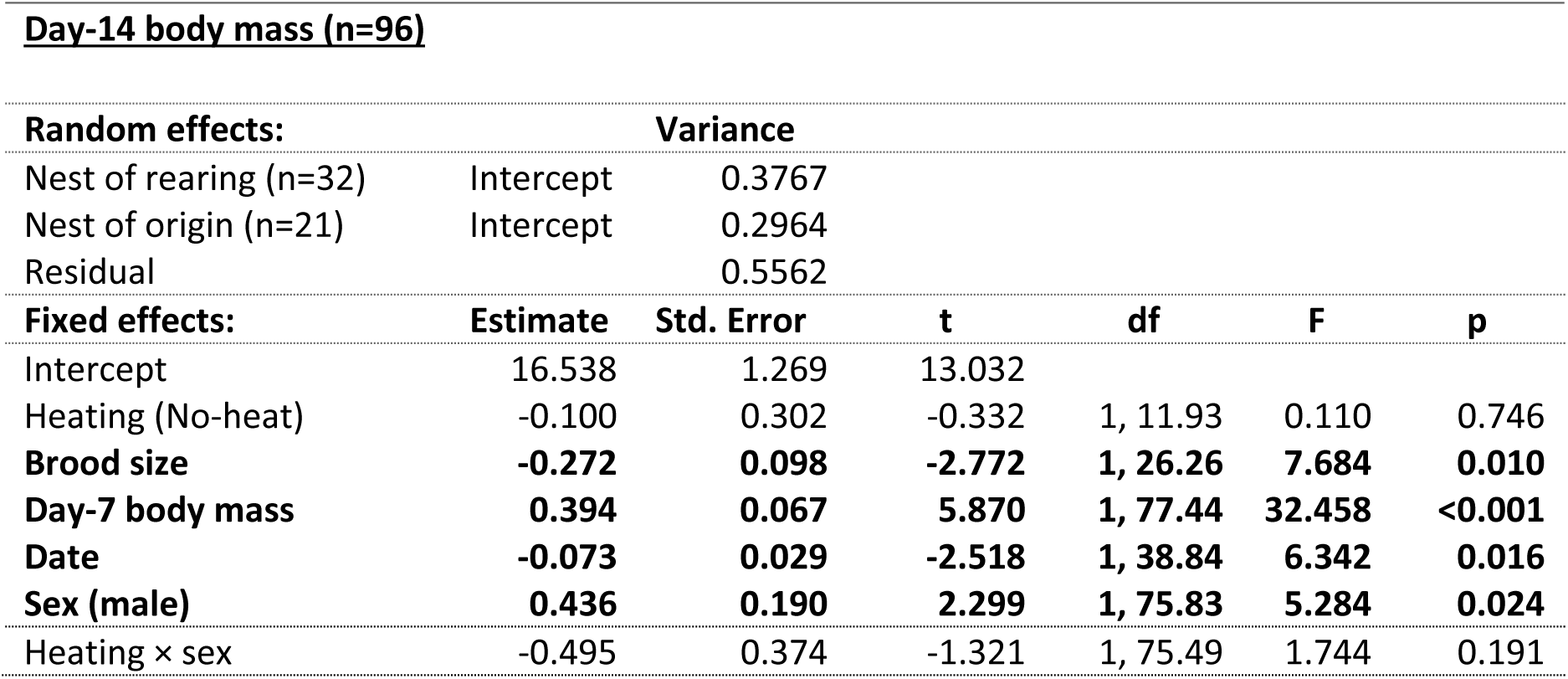
Results of linear mixed models for nestling body mass.

| **Day-14 body mass (n=96)** |  |  |  |  |  |  |
| --- | --- | --- | --- | --- | --- | --- |
| **Random effects:** |  | **Variance** |  |  |  |  |
| Nest of rearing (n=32) | Intercept | 0.3767 |  |  |  |  |
| Nest of origin (n=21) | Intercept | 0.2964 |  |  |  |  |
| Residual |  | 0.5562 |  |  |  |  |
| **Fixed effects:** | **Estimate** | **Std. Error** | **t** | **df** | **F** | **p** |
| Intercept | 16.538 | 1.269 | 13.032 |  |  |  |
| Heating (No-heat) | -0.100 | 0.302 | -0.332 | 1, 11.93 | 0.110 | 0.746 |
| **Brood size** | **-0.272** | **0.098** | **-2.772** | **1, 26.26** | **7.684** | **0.010** |
| **Day-7 body mass** | **0.394** | **0.067** | **5.870** | **1, 77.44** | **32.458** | **<0.001** |
| **Date** | **-0.073** | **0.029** | **-2.518** | **1, 38.84** | **6.342** | **0.016** |
| **Sex (male)** | **0.436** | **0.190** | **2.299** | **1, 75.83** | **5.284** | **0.024** |
| Heating × sex | -0.495 | 0.374 | -1.321 | 1, 75.49 | 1.744 | 0.191 |

**Table S3:**
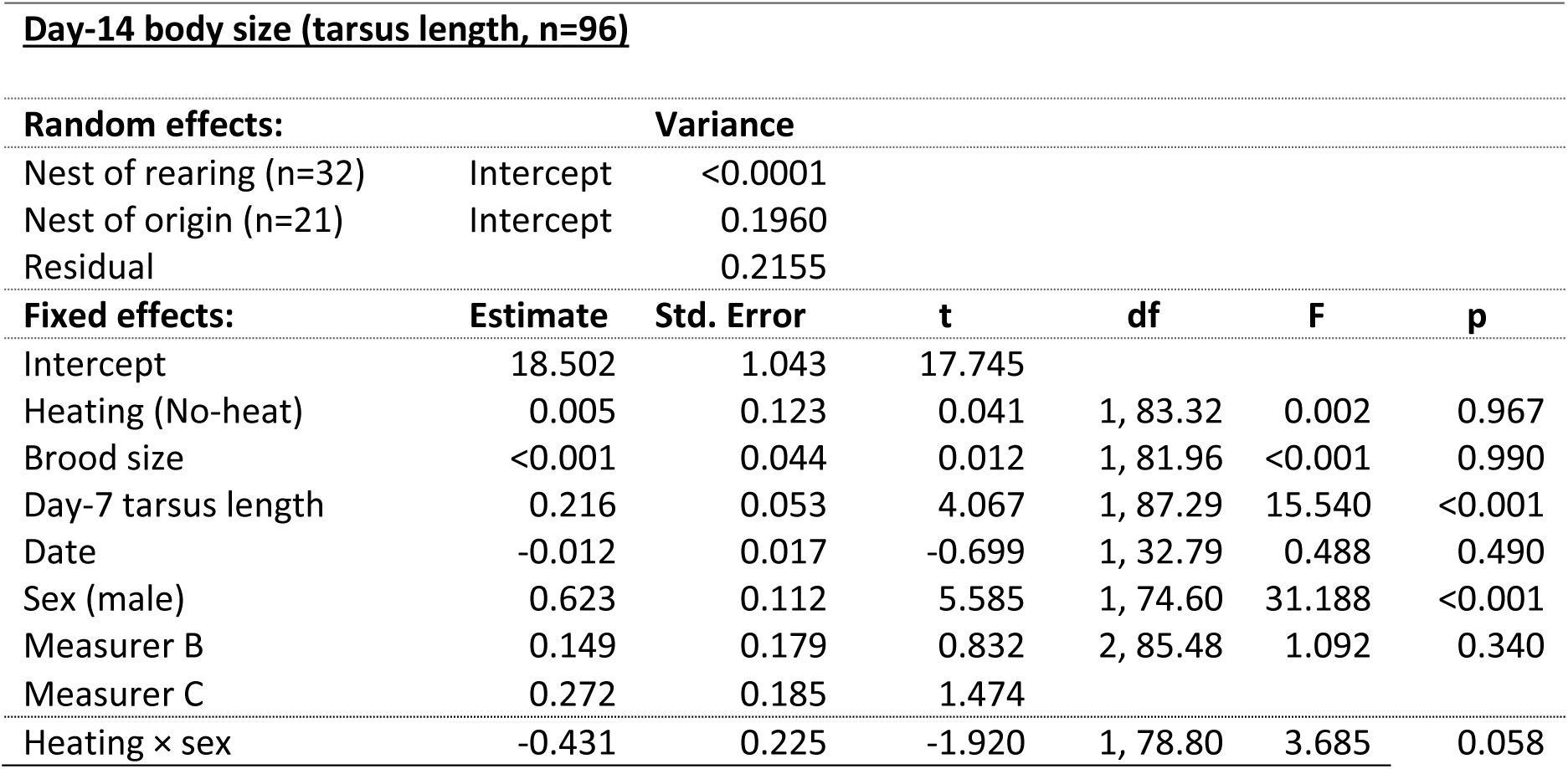
Results of linear mixed models for nestling body size.

| **Day-14 body size (tarsus length, n=96)** |  |  |  |  |  |  |
| --- | --- | --- | --- | --- | --- | --- |
| **Random effects:** |  | **Variance** |  |  |  |  |
| Nest of rearing (n=32) | Intercept | <0.0001 |  |  |  |  |
| Nest of origin (n=21) | Intercept | 0.1960 |  |  |  |  |
| Residual |  | 0.2155 |  |  |  |  |
| **Fixed effects:** | **Estimate** | **Std. Error** | **t** | **df** | **F** | **p** |
| Intercept | 18.502 | 1.043 | 17.745 |  |  |  |
| Heating (No-heat) | 0.005 | 0.123 | 0.041 | 1, 83.32 | 0.002 | 0.967 |
| Brood size | <0.001 | 0.044 | 0.012 | 1, 81.96 | <0.001 | 0.990 |
| Day-7 tarsus length | 0.216 | 0.053 | 4.067 | 1, 87.29 | 15.540 | <0.001 |
| Date | -0.012 | 0.017 | -0.699 | 1, 32.79 | 0.488 | 0.490 |
| Sex (male) | 0.623 | 0.112 | 5.585 | 1, 74.60 | 31.188 | <0.001 |
| Measurer B | 0.149 | 0.179 | 0.832 | 2, 85.48 | 1.092 | 0.340 |
| Measurer C | 0.272 | 0.185 | 1.474 |  |  |  |
| Heating × sex | -0.431 | 0.225 | -1.920 | 1, 78.80 | 3.685 | 0.058 |

**Table S4:** Results of linear mixed models for juvenile body mass.

| **Juvenile body mass (n=25)** |  |  |  |  |  |  |
| --- | --- | --- | --- | --- | --- | --- |
| **Random effects:** |  | **Variance** |  |  |  |  |
| Nest of rearing (n=16) | Intercept | 0.0125 |  |  |  |  |
| Nest of origin (n=13) | Intercept | <0.0001 |  |  |  |  |
| Residual |  | 0.8796 |  |  |  |  |
| **Fixed effects:** | **Estimate** | **Std. Error** | **t** | **df** | **F** | **p** |
| Intercept | 13.911 | 3.985 | 3.491 |  |  |  |
| **Heating (No-heat)** | **-1.057** | **0.418** | **-2.532** | **1, 13.97** | **6.413** | **0.024** |
| Brood size | 0.141 | 0.134 | 1.051 | 1, 14.39 | 1.105 | 0.311 |
| Sex (male) | 0.408 | 0.419 | 0.973 | 1, 19.02 | 0.946 | 0.343 |
| Day-14 body mass | 0.225 | 0.204 | 1.104 | 1, 19.96 | 1.219 | 0.283 |
| Heating × sex | 0.051 | 0.956 | 0.053 | 1, 19.00 | 0.003 | 0.958 |

**Table S5:**
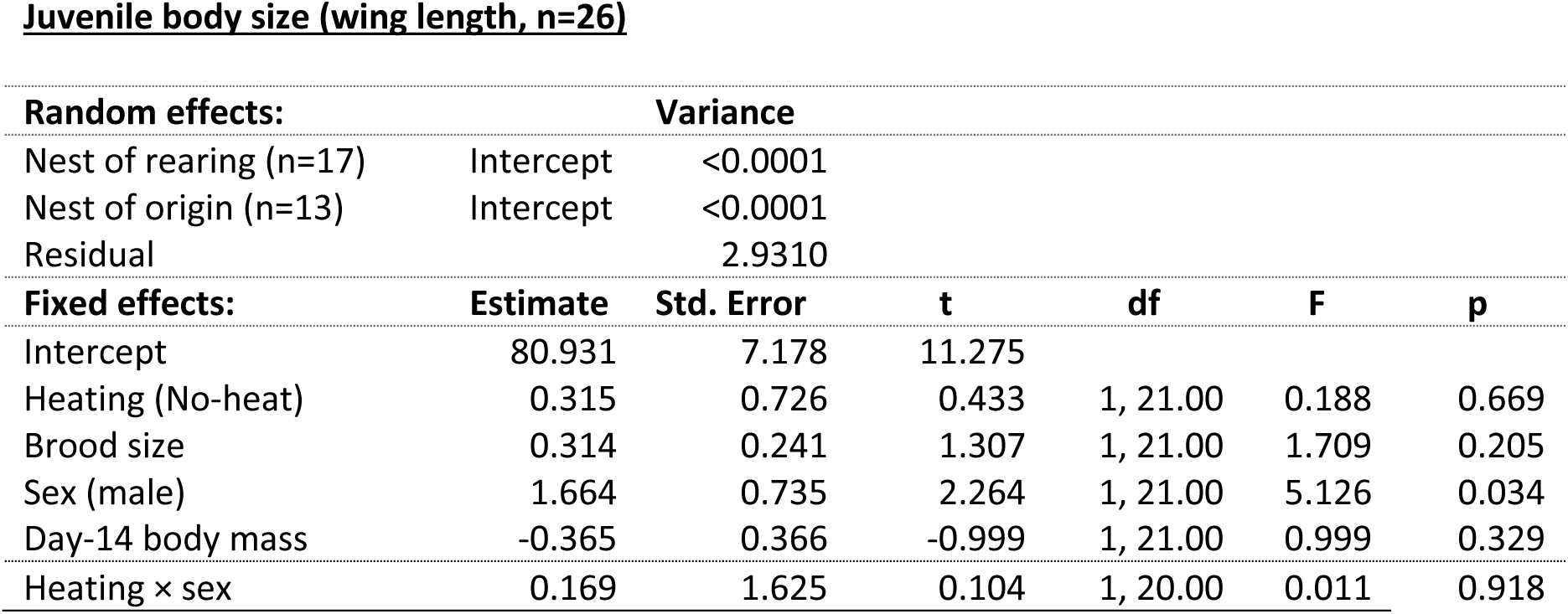
Results of linear mixed models for juvenile body size.

**Juvenile body size (wing length, n=26)**
| **Random effects:** |  | **Variance** |  |  |  |  |
| --- | --- | --- | --- | --- | --- | --- |
| Nest of rearing (n=17) | Intercept | <0.0001 |  |  |  |  |
| Nest of origin (n=13) | Intercept | <0.0001 |  |  |  |  |
| Residual |  | 2.9310 |  |  |  |  |
| **Fixed effects:** | **Estimate** | **Std. Error** | **t** | **df** | **F** | **p** |
| Intercept | 80.931 | 7.178 | 11.275 |  |  |  |
| Heating (No-heat) | 0.315 | 0.726 | 0.433 | 1, 21.00 | 0.188 | 0.669 |
| Brood size | 0.314 | 0.241 | 1.307 | 1, 21.00 | 1.709 | 0.205 |
| Sex (male) | 1.664 | 0.735 | 2.264 | 1, 21.00 | 5.126 | 0.034 |
| Day-14 body mass | -0.365 | 0.366 | -0.999 | 1, 21.00 | 0.999 | 0.329 |
| Heating × sex | 0.169 | 1.625 | 0.104 | 1, 20.00 | 0.011 | 0.918 |

**Table S6:**
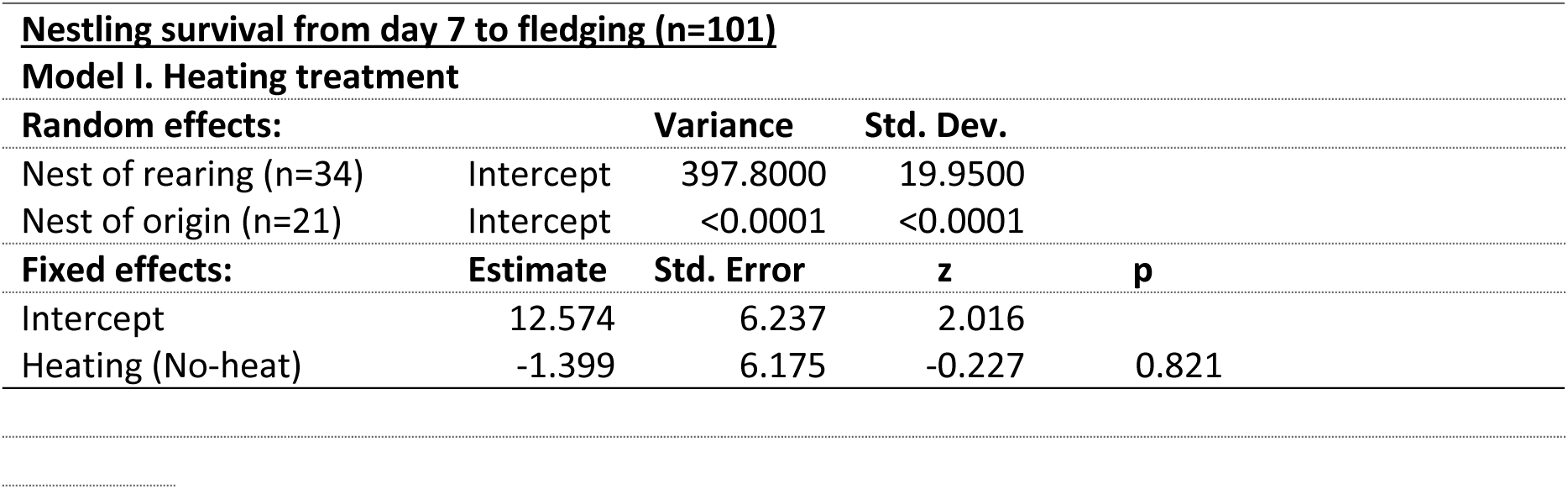
Results of generalized linear mixed models for nestling survival. For survival, binomial GLMMs were fitted with maximum likelihood by Laplace approximation. As there were only 7 nestlings that failed to fledge, other covariates (sex and day-7 body mass) were excluded in these models to enable model convergence.

**Table S7:** Results of generalized linear mixed models for juvenile survival.

| <u>Post-fledging survival (n=97)</u> |  |  |  |  |  |
| --- | --- | --- | --- | --- | --- |
| Random effects: |  | Variance | Std. Dev. |  |  |
| Nest of rearing (n=32) | Intercept | 1.0590 | 1.0293 |  |  |
| Nest of origin (n=21) | Intercept | <0.0001 | 0.0014 |  |  |
| Fixed effects: |  | Estimate | Std. Error | z | p |
| Intercept |  | -9.411 | 4.840 | -1.944 |  |
| Heating (No-heat) |  | 0.837 | 0.687 | 1.218 | 0.223 |
| Day-14 body mass |  | 0.399 | 0.257 | 1.553 | 0.120 |
| Sex (male) |  | 0.812 | 0.629 | 1.291 | 0.197 |
| Heating × sex |  | -0.174 | 1.157 | -0.151 | 0.880 |

**Table S8:**
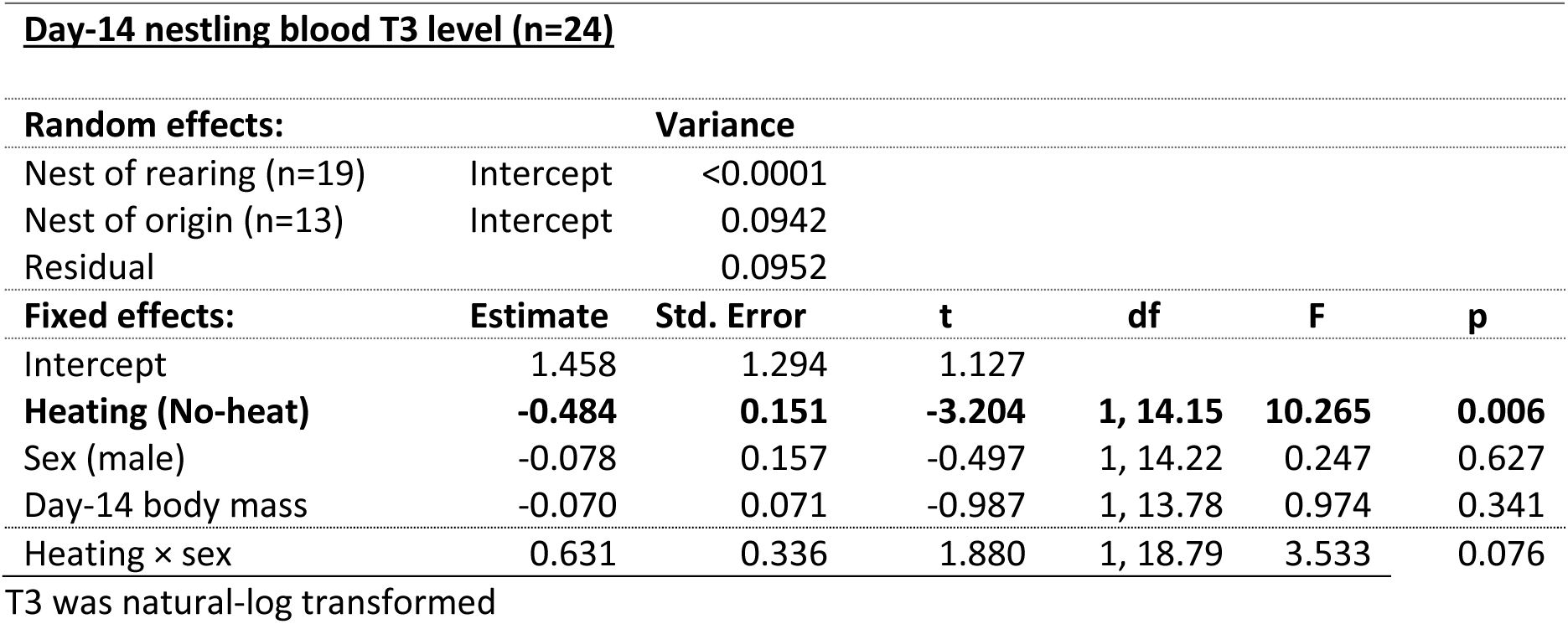
Results of linear mixed models for nestling plasma T3 levels.

| **Day-14 nestling blood T3 level (n=24)** |  |  |  |  |  |  |
| --- | --- | --- | --- | --- | --- | --- |
| **Random effects:** |  | **Variance** |  |  |  |  |
| Nest of rearing (n=19) | Intercept | <0.0001 |  |  |  |  |
| Nest of origin (n=13) | Intercept | 0.0942 |  |  |  |  |
| Residual |  | 0.0952 |  |  |  |  |
| **Fixed effects:** | **Estimate** | **Std. Error** | **t** | **df** | **F** | **p** |
| Intercept | 1.458 | 1.294 | 1.127 |  |  |  |
| **Heating (No-heat)** | **-0.484** | **0.151** | **-3.204** | **1, 14.15** | **10.265** | **0.006** |
| Sex (male) | -0.078 | 0.157 | -0.497 | 1, 14.22 | 0.247 | 0.627 |
| Day-14 body mass | -0.070 | 0.071 | -0.987 | 1, 13.78 | 0.974 | 0.341 |
| Heating × sex | 0.631 | 0.336 | 1.880 | 1, 18.79 | 3.533 | 0.076 |

T3 was natural-log transformed

**Table S9:** Results of linear mixed models for nestling plasma T4 levels.

| **Day-14 nestling blood T4 level (n=24)** |  |  |  |  |  |  |
| --- | --- | --- | --- | --- | --- | --- |
| **Random effects:** |  | **Variance** |  |  |  |  |
| Nest of rearing (n=19) | Intercept | <0.0001 |  |  |  |  |
| Nest of origin (n=13) | Intercept | 0.0934 |  |  |  |  |
| Residual |  | 0.0525 |  |  |  |  |
| **Fixed effects:** | **Estimate** | **Std. Error** | **t** | **df** | **F** | **p** |
| Intercept | 2.423 | 1.004 | 2.413 |  |  |  |
| **Heating (No-heat)** | **-0.283** | **0.118** | **-2.403** | **1, 14.69** | **5.776** | **0.030** |
| Sex (male) | 0.233 | 0.122 | 1.905 | 1, 14.68 | 3.627 | 0.077 |
| Day-14 body mass | -0.033 | 0.055 | -0.599 | 1, 14.10 | 0.358 | 0.559 |
| Heating × sex | 0.293 | 0.288 | 1.019 | 1, 17.80 | 1.038 | 0.322 |

T4 was natural-log transformed.

**Table S10:** Results of linear mixed models for nestling blood oxidative damage to lipids (MDA) levels.

| **Day-14 nestling blood MDA level (n=57)** |  |  |  |  |  |  |
| --- | --- | --- | --- | --- | --- | --- |
| **Random effects:** |  | **Variance** |  |  |  |  |
| Nest of rearing (n=26) | Intercept | 0.0005 |  |  |  |  |
| Nest of origin (n=18) | Intercept | <0.0001 |  |  |  |  |
| Batch of TBARS (n=6) | Intercept | 0.0994 |  |  |  |  |
| Residual |  | 0.0519 |  |  |  |  |
| **Fixed effects:** | **Estimate** | **Std. Error** | **t** | **df** | **F** | **p** |
| Intercept | -2.991 | 0.530 | -5.642 |  |  |  |
| Heating (No-heat) | 0.033 | 0.074 | 0.447 | 1, 22.68 | 0.200 | 0.659 |
| Sex (male) | 0.017 | 0.071 | 0.240 | 1, 47.99 | 0.058 | 0.811 |
| Day-14 body mass | -0.018 | 0.028 | -0.629 | 1, 37.33 | 0.396 | 0.533 |
| Heating × sex | 0.027 | 0.146 | 0.183 | 1, 47.53 | 0.034 | 0.855 |

**Table S11:** Results of linear mixed models for nestling blood total glutathione (tGSH) levels.

| **Day-14 nestling blood tGSH level (n=56)** |  |  |  |  |  |  |
| --- | --- | --- | --- | --- | --- | --- |
| **Random effects:** |  | **Variance** |  |  |  |  |
| Nest of rearing (n=27) | Intercept | <0.0001 |  |  |  |  |
| Nest of origin (n=18) | Intercept | 0.0237 |  |  |  |  |
| Batch of GSH (n=2) | Intercept | 0.0208 |  |  |  |  |
| Residual |  | 0.0936 |  |  |  |  |
| **Fixed effects:** | **Estimate** | **Std. Error** | **t** | **df** | **F** | **p** |
| Intercept | -0.226 | 0.773 | -0.292 |  |  |  |
| Heating (No-heat) | -0.098 | 0.095 | -1.031 | 1, 49.86 | 1.062 | 0.308 |
| Sex (male) | -0.007 | 0.097 | -0.067 | 1, 50.20 | 0.005 | 0.947 |
| Day-14 body mass | -0.056 | 0.042 | -1.351 | 1, 49.76 | 1.824 | 0.183 |
| Heating × sex | 0.247 | 0.194 | 1.278 | 1, 47.89 | 1.634 | 0.208 |

**Table S12:**
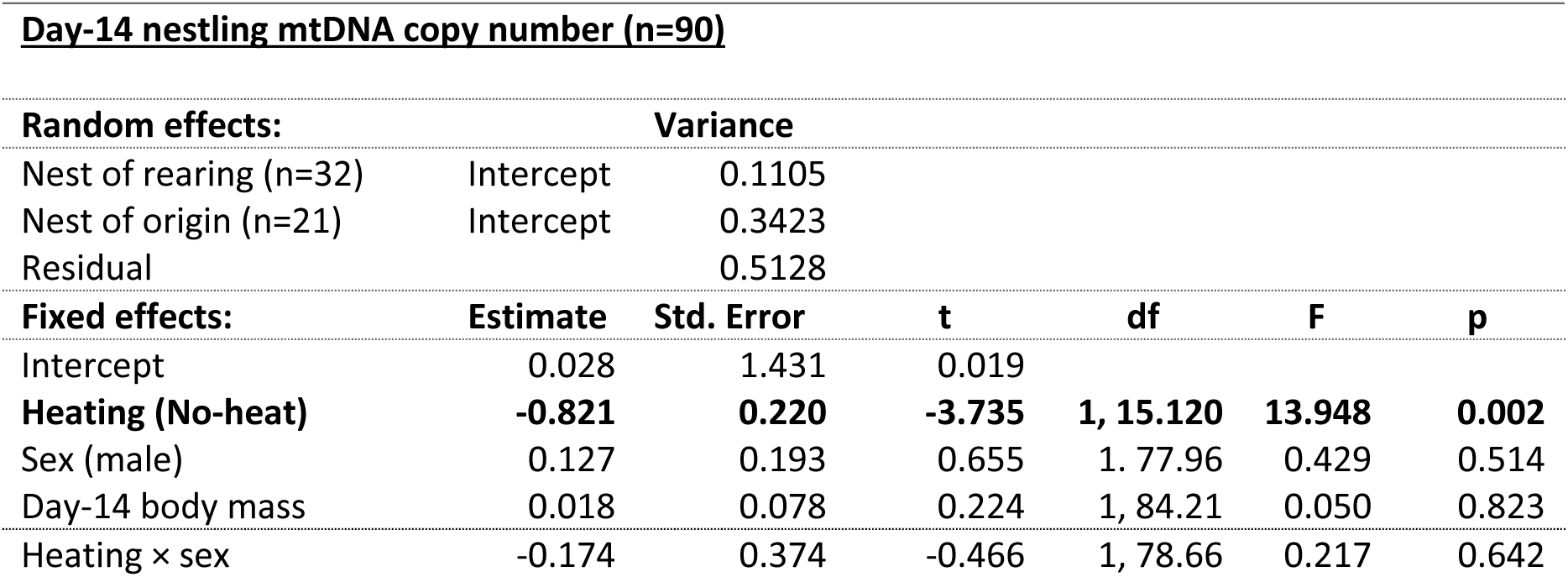
Results of linear mixed models for nestling blood cell mtDNA copy number.

**Table S13:** Results of linear mixed models for juvenile blood cell mtDNA copy number.

| **Juvenile mtDNA copy number (n=23)** |  |  |  |  |  |  |
| --- | --- | --- | --- | --- | --- | --- |
| **Random effects:** |  | **Variance** |  |  |  |  |
| Nest of rearing (n=14) | Intercept | <0.0001 |  |  |  |  |
| Nest of origin (n=13) | Intercept | <0.0001 |  |  |  |  |
| Residual |  | 0.9976 |  |  |  |  |
| **Fixed effects:** | **Estimate** | **Std. Error** | **t** | **df** | **F** | **p** |
| Intercept | -7.514 | 4.240 | -1.772 |  |  |  |
| Heating (No-heat) | 0.404 | 0.515 | 0.784 | 1, 19.00 | 0.615 | 0.443 |
| Sex (male) | -0.237 | 0.441 | -0.538 | 1, 19.00 | 0.289 | 0.597 |
| Juvenile body mass | 0.404 | 0.224 | 1.803 | 1, 19.00 | 3.250 | 0.087 |
| Heating × sex | 0.833 | 0.936 | 0.891 | 1, 17.45 | 0.794 | 0.385 |

**Table S14:** Results of linear mixed models for nestling blood cell relative telomere length.

| **Day-14 nestling telomere length (n=90)** |  |  |  |  |  |  |
| --- | --- | --- | --- | --- | --- | --- |
| **Random effects:** |  | **Variance** |  |  |  |  |
| Nest of rearing (n=32) | Intercept | 0.3542 |  |  |  |  |
| Nest of origin (n=21) | Intercept | 0.0614 |  |  |  |  |
| Residual |  | 0.6041 |  |  |  |  |
| **Fixed effects:** | **Estimate** | **Std. Error** | **t** | **df** | **F** | **p** |
| Intercept | -1.977 | 1.534 | -1.288 |  |  |  |
| Heating (No-heat) | 0.161 | 0.280 | 0.574 | 1, 15.83 | 0.330 | 0.574 |
| Sex (male) | -0.212 | 0.210 | -1.011 | 1, 81.66 | 1.022 | 0.315 |
| Day-14 body mass | 0.110 | 0.084 | 1.317 | 1, 84.37 | 1.735 | 0.191 |
| Heating × sex | -0.264 | 0.399 | -0.663 | 1, 76.04 | 0.439 | 0.510 |

**Table S15:**
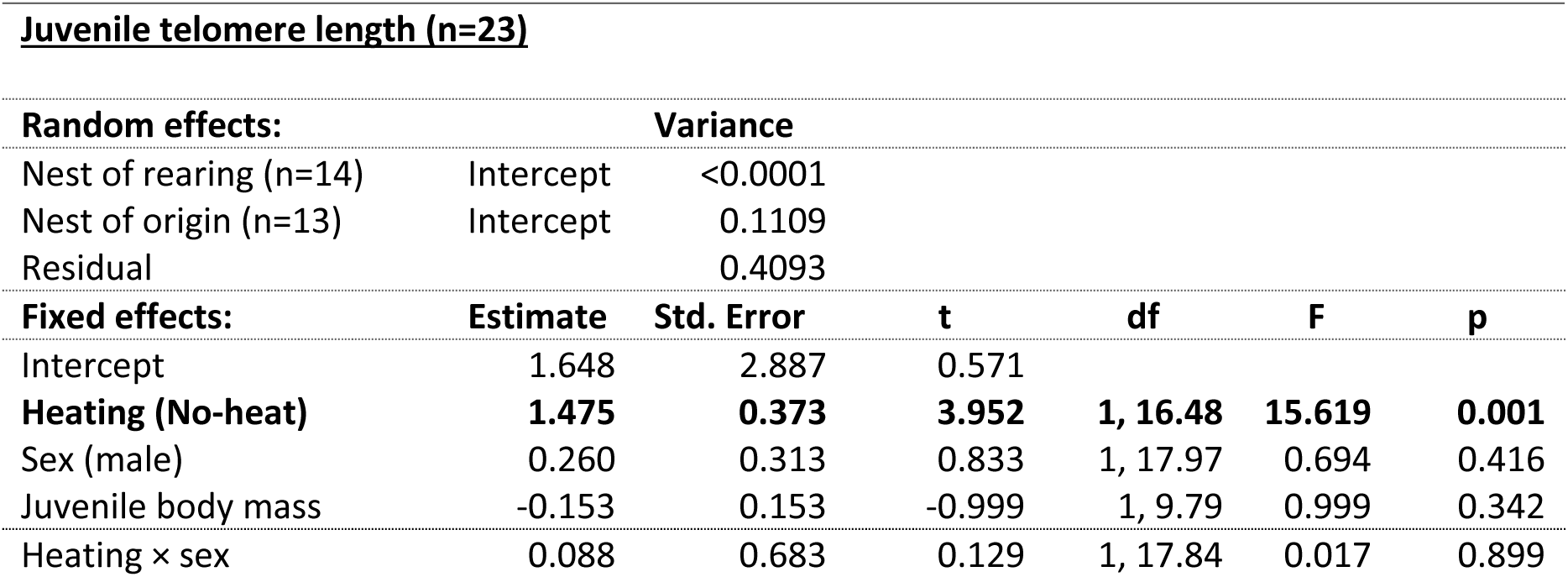
Results of linear mixed models for juvenile blood cell relative telomere length.

**Table S16:**
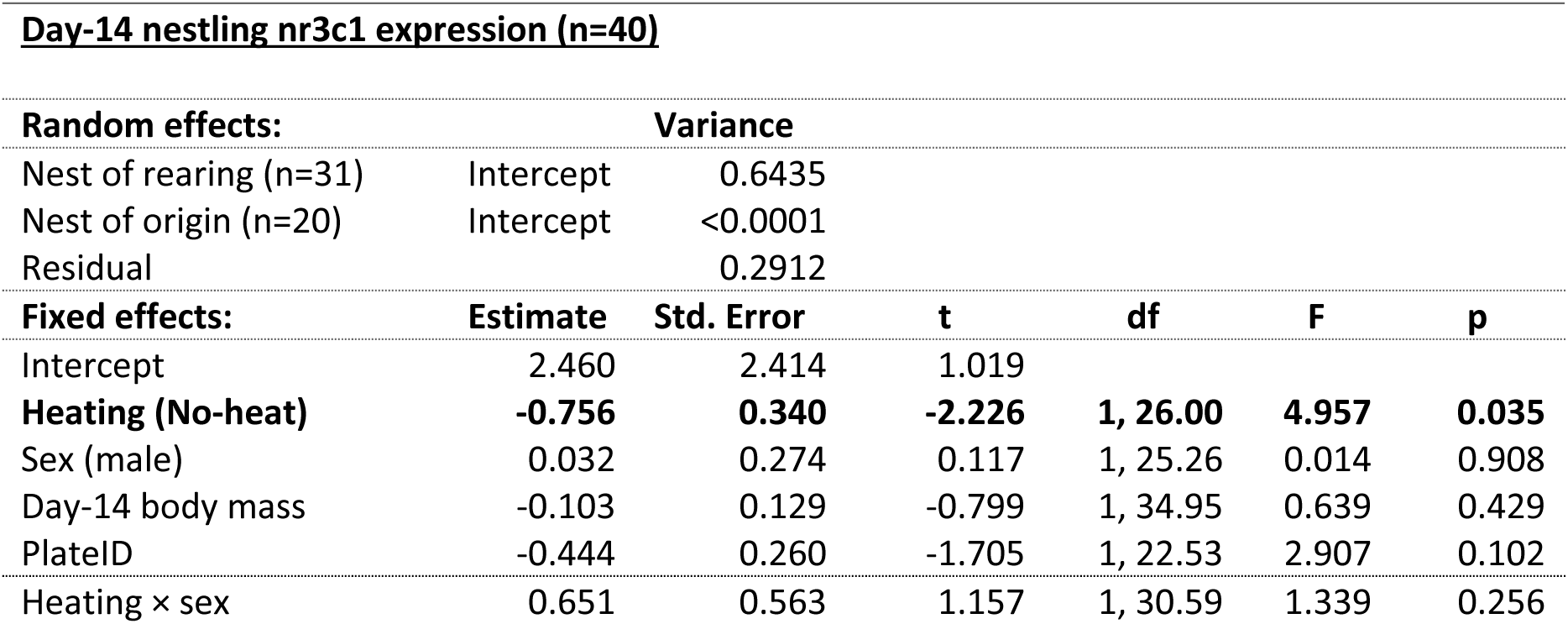
Results of linear mixed models for nr3c1 expression levels.

**Table S17:**
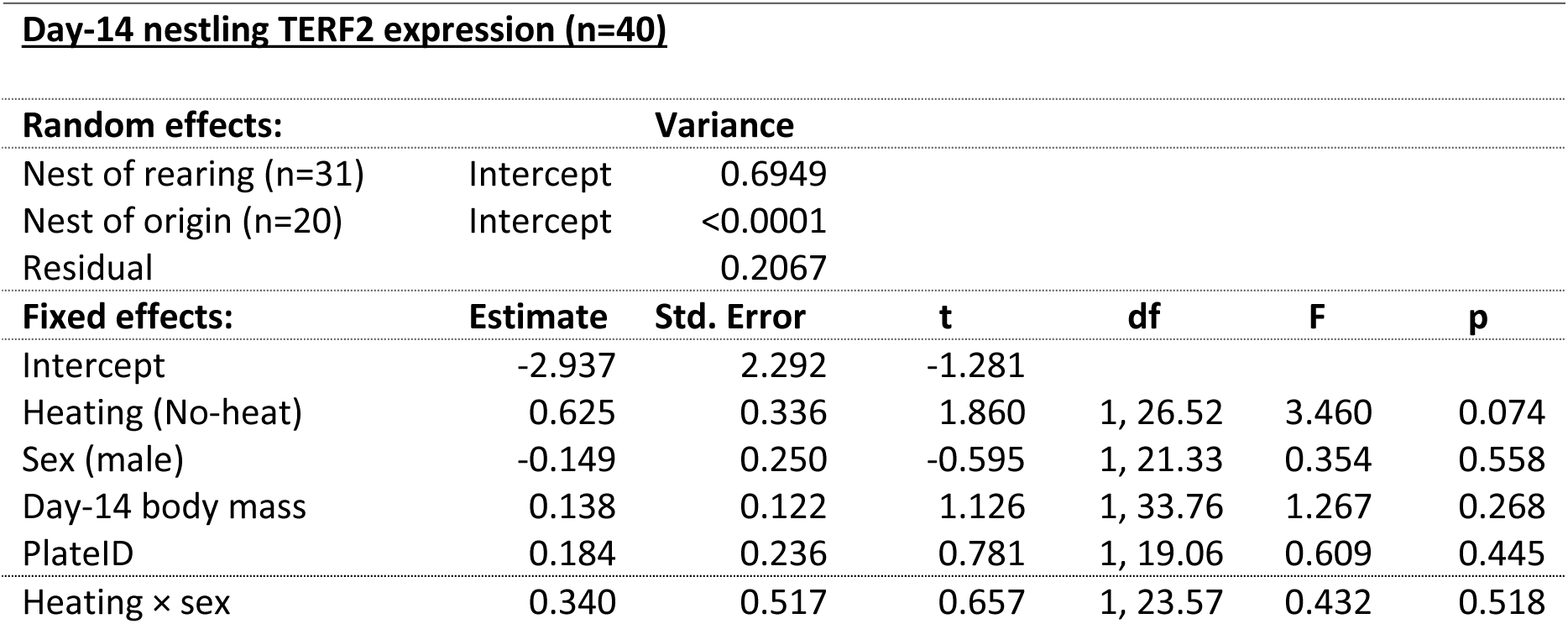
Results of linear mixed models for TERF2 expression levels.

| **Day-14 nestling TERF2 expression (n=40)** |  |  |  |  |  |  |
| --- | --- | --- | --- | --- | --- | --- |
| **Random effects:** |  | **Variance** |  |  |  |  |
| Nest of rearing (n=31) | Intercept | 0.6949 |  |  |  |  |
| Nest of origin (n=20) | Intercept | <0.0001 |  |  |  |  |
| Residual |  | 0.2067 |  |  |  |  |
| **Fixed effects:** | **Estimate** | **Std. Error** | **t** | **df** | **F** | **p** |
| Intercept | -2.937 | 2.292 | -1.281 |  |  |  |
| Heating (No-heat) | 0.625 | 0.336 | 1.860 | 1, 26.52 | 3.460 | 0.074 |
| Sex (male) | -0.149 | 0.250 | -0.595 | 1, 21.33 | 0.354 | 0.558 |
| Day-14 body mass | 0.138 | 0.122 | 1.126 | 1, 33.76 | 1.267 | 0.268 |
| PlateID | 0.184 | 0.236 | 0.781 | 1, 19.06 | 0.609 | 0.445 |
| Heating × sex | 0.340 | 0.517 | 0.657 | 1, 23.57 | 0.432 | 0.518 |

**Table S18:** Results of linear mixed models for TERT expression levels.

| **Day-14 nestling TERT expression (n=40)** |  |  |  |  |  |  |
| --- | --- | --- | --- | --- | --- | --- |
| **Random effects:** |  | **Variance** |  |  |  |  |
| Nest of rearing (n=31) | Intercept | <0.0001 |  |  |  |  |
| Nest of origin (n=20) | Intercept | 0.4313 |  |  |  |  |
| Residual |  | 0.5662 |  |  |  |  |
| **Fixed effects:** | **Estimate** | **Std. Error** | **t** | **df** | **F** | **p** |
| Intercept | -1.171 | 2.426 | -0.483 |  |  |  |
| Heating (No-heat) | 0.537 | 0.274 | 1.959 | 1, 27.07 | 3.837 | 0.060 |
| Sex (male) | -0.116 | 0.293 | -0.397 | 1, 30.38 | 0.158 | 0.694 |
| Day-14 body mass | 0.051 | 0.131 | 0.389 | 1, 34.85 | 0.151 | 0.700 |
| PlateID | 0.039 | 0.278 | 0.141 | 1, 27.25 | 0.020 | 0.889 |
| Heating × sex | 0.070 | 0.621 | 0.113 | 1, 32.79 | 0.013 | 0.320 |

**Table S19:** Results of linear mixed models for HSP70 expression levels.

| **Day-14 nestling HSP70 expression (n=40)** |  |  |  |  |  |  |
| --- | --- | --- | --- | --- | --- | --- |
| **Random effects:** |  | **Variance** |  |  |  |  |
| Nest of rearing (n=31) | Intercept | 0.7109 |  |  |  |  |
| Nest of origin (n=20) | Intercept | <0.0001 |  |  |  |  |
| Residual |  | 0.2268 |  |  |  |  |
| **Fixed effects:** | **Estimate** | **Std. Error** | **t** | **df** | **F** | **p** |
| Intercept | -2.537 | 2.353 | -1.078 |  |  |  |
| Heating (No-heat) | 0.378 | 0.342 | 1.104 | 1, 27.26 | 1.219 | 0.279 |
| Sex (male) | -0.008 | 0.259 | -0.033 | 1, 23.05 | 0.001 | 0.974 |
| Day-14 body mass | 0.119 | 0.125 | 0.950 | 1, 34.19 | 0.902 | 0.349 |
| PlateID | 0.183 | 0.244 | 0.749 | 1, 20.76 | 0.561 | 0.462 |
| Heating × sex | 0.022 | 0.539 | 0.040 | 1, 26.08 | 0.002 | 0.968 |

**Table S20:** Results of linear mixed models for HSP90 expression levels.

| **Day-14 nestling HSP90 expression (n=40)** |  |  |  |  |  |  |
| --- | --- | --- | --- | --- | --- | --- |
| **Random effects:** |  | **Variance** |  |  |  |  |
| Nest of rearing (n=31) | Intercept | 0.4735 |  |  |  |  |
| Nest of origin (n=20) | Intercept | <0.0001 |  |  |  |  |
| Residual |  | 0.3562 |  |  |  |  |
| **Fixed effects:** | **Estimate** | **Std. Error** | **t** | **df** | **F** | **p** |
| Intercept | -7.956 | 2.331 | -3.414 |  |  |  |
| Heating (No-heat) | -0.006 | 0.317 | -0.019 | 1, 25.35 | <0.001 | 0.985 |
| Sex (male) | -0.153 | 0.275 | -0.558 | 1, 29.55 | 0.311 | 0.581 |
| **Day-14 body mass** | **0.419** | **0.125** | **3.360** | **1, 34.56** | **11.290** | **0.002** |
| Plate | 0.308 | 0.263 | 1.170 | 1, 26.56 | 1.370 | 0.252 |
| Heating × sex | 0.1885 | 0.557 | 0.338 | 1, 31.31 | 0.115 | 0.734 |

**Table S21:** Results of linear mixed models for NRF2 expression levels.

| **Day-14 nestling NRF2 expression (n=40)** |  |  |  |  |  |  |
| --- | --- | --- | --- | --- | --- | --- |
| **Random effects:** |  | **Variance** |  |  |  |  |
| Nest of rearing (n=31) | Intercept | 0.4850 |  |  |  |  |
| Nest of origin (n=20) | Intercept | <0.0001 |  |  |  |  |
| Residual |  | 0.4215 |  |  |  |  |
| **Fixed effects:** | **Estimate** | **Std. Error** | **t** | **df** | **F** | **p** |
| Intercept | -6.242 | 2.446 | -2.552 |  |  |  |
| Heating (No-heat) | 0.233 | 0.330 | 0.705 | 1, 26.04 | 0.497 | 0.487 |
| Sex (male) | 0.294 | 0.291 | 1.011 | 1, 31.03 | 1.022 | 0.320 |
| **Day-14 body mass** | **0.308** | **0.131** | **2.354** | **1, 34.33** | **5.541** | **0.024** |
| Plate | 0.316 | 0.279 | 1.131 | 1, 28.30 | 1.278 | 0.268 |
| Heating × sex | -0.047 | 0.591 | -0.080 | 1, 32.64 | 0.006 | 0.937 |

## Notes

https://doi.org/10.6084/m9.figshare.28838297.v1

### Competing Interest Statement

The authors have declared no competing interest.

### Summary of Updates

Figures have been changed to be more readable, and methodological details were added

## References

1. McKechnie AE, Rushworth IA, Myburgh F, Cunningham SJ. Mortality among birds and bats during an extreme heat event in eastern South Africa. Austral Ecology. 2021. doi:10.1111/aec.13025

2. Conradie SR, Woodborne SM, Cunningham SJ, McKechnie AE. Chronic, sublethal effects of high temperatures will cause severe declines in southern African arid-zone birds during the 21st century. Proceedings Of The National Academy Of Sciences Of The United States Of America. 2019;116: 14065–14070. doi:10.1073/pnas.1821312116

3. Boyles JG, Seebacher F, Smit B, McKechnie AE. Adaptive Thermoregulation in Endotherms May Alter Responses to Climate Change. Integrative and Comparative Biology. 2011;51: 676–690. doi:10.1093/icb/icr053

4. Diehl JN, Alton LA, White CR, Peters A. Thermoregulatory strategies of songbird nestlings reveal limited capacity for cooling and high risk of dehydration. J Therm Biol. 2023;117: 103707. doi:10.1016/j.jtherbio.2023.103707

5. Metcalfe N, Monaghan P. Compensation for a bad start: grow now, pay later? Trends in Ecology & Evolution. 2001;16: 254–260. Available:

6. Preston JD, Reynolds LJ, Pearson KJ. Developmental Origins of Health Span and Life Span: A Mini-Review. Gerontology. 2018;64: 237–245. doi:10.1159/000485506

7. Greño JL, Belda EJ, Barba E. Influence of temperatures during the nestling period on post-fledging survival of great tit Parus major in a Mediterranean habitat. J Avian Biol. 2008;39: 41–49. doi:10.1111/j.0908-8857.2008.04120.x

8. Hepp GR, Kennamer RA. Warm is better: incubation temperature influences apparent survival and recruitment of wood ducks (Aix sponsa). PLoS ONE. 2012;7: e47777. doi:10.1371/journal.pone.0047777

9. Berntsen HH, Bech C. Incubation temperature influences survival in a small passerine bird. Journal of Avian Biology. 2015; n/a-n/a. doi:10.1111/jav.00688

10. Nord A, Nilsson J-A. Long-term consequences of high incubation temperature in a wild bird population. Biology Letters. 2016; 1–4. doi:10.1098/rsbl.2016.0087

11. Andreasson F, Nord A, Nilsson J-A. Experimentally increased nest temperature affects body temperature, growth and apparent survival in blue tit nestlings. Journal of Avian Biology. 2018; 1–14. doi:10.1111/jav

12. Catalano R, Bruckner T, Smith KR. Ambient temperature predicts sex ratios and male longevity. Proc National Acad Sci. 2008;105: 2244–2247. doi:10.1073/pnas.0710711104

13. Bowen SJ, Washburn KW. Thyroid and Adrenal Response to Heat Stress in Chickens and Quail Differing in Heat Tolerance1. Poultry science. 1985;64: 149–154. doi:10.3382/ps.0640149

14. Iqbal A, Decuypere E, Azim AAE. Pre-and post-hatch high temperature exposure affects the thyroid hormones and corticosterone response to acute heat stress in growing chicken (Gallus domesticus). Journal of Thermal Biology. 1990;15: 149–153. doi:10.1016/0306-4565(90)90032-d

15. Salgado RM, Sheard AC, Vaughan RA, Parker DL, Schneider SM, Kenefick RW, et al. Mitochondrial efficiency and exercise economy following heat stress: a potential role of uncoupling protein 3. Physiological Reports. 2017;5: e13054. doi:10.14814/phy2.13054

16. Kikusato M, Ramsey JJ, Amo T, Toyomizu M. Application of modular kinetic analysis to mitochondrial oxidative phosphorylation in skeletal muscle of birds exposed to acute heat stress. FEBS Letters. 2010;584: 3143–3148. doi:10.1016/j.febslet.2010.05.057

17. Balaban RS, Nemoto S, Finkel T. Mitochondria, Oxidants, and Aging. Cell. 2005;120: 483–495. doi:10.1016/j.cell.2005.02.001

18. Dawson RD, Lawrie CC, O’Brien EL. The importance of microclimate variation in determining size, growth and survival of avian offspring: experimental evidence from a cavity nesting passerine. Oecologia. 2005;144: 499–507. doi:10.1007/s00442-005-0075-7

19. Rodríguez S, Barba E. Nestling Growth is Impaired by Heat Stress: an Experimental Study in a Mediterranean Great Tit Population. Zoological studies. 2016;55: e40. doi:10.6620/zs.2016.55-40

20. Wilbourn RV, Moatt JP, Froy H, Walling CA, Nussey DH, Boonekamp JJ. The relationship between telomere length and mortality risk in non-model vertebrate systems: a meta-analysis. Philosophical transactions of the Royal Society of London Series B, Biological sciences. 2018;373: 20160447–9. doi:10.1098/rstb.2016.0447

21. Eastwood JR, Hall ML, Teunissen N, Kingma SA, Aranzamendi NH, Fan M, et al. Early-life telomere length predicts lifespan and lifetime reproductive success in a wild bird. Molecular Ecology. 2019;28: 1127–1137. doi:10.1111/mec.15002

22. Stier A, Metcalfe NB, Monaghan P. Pace and stability of embryonic development affect telomere dynamics: an experimental study in a precocial bird model. Proceedings of the Royal Society of London Series B: Biological Sciences. 2020;287: 20201378–9. doi:10.1098/rspb.2020.1378

23. Chatelain M, Drobniak SM, Szulkin M. The association between stressors and telomeres in non-human vertebrates: a meta-analysis. Cote J, editor. Ecology Letters. 2019;99: 21–18. doi:10.1111/ele.13426

24. Reichert S, Stier A. Does oxidative stress shorten telomeres in vivo? A review. Biology Letters. 2017;13: 20170463–7. doi:10.1098/rsbl.2017.0463

25. Liu L, Trimarchi JR, Smith PJS, Keefe DL. Mitochondrial dysfunction leads to telomere attrition and genomic instability. Aging Cell. 2002;1: 40–46. Available:

26. Casagrande S, Hau M. Telomere attrition: metabolic regulation and signalling function? Biology Letters. 2019;15: 20180885–11. doi:10.1098/rsbl.2018.0885

27. Simide R, Angelier F, Gaillard S, Stier A. Age and Heat Stress as Determinants of Telomere Length in a Long-Lived Fish, the Siberian Sturgeon. Physiological And Biochemical Zoology. 2016;89: 441–447. doi:10.1086/687378

28. Debes PV, Visse M, Panda B, Ilmonen P, Vasemägi A. Is telomere length a molecular marker of past thermal stress in wild fish? Molecular Ecology. 2016. doi:10.1111/mec.13856

29. Zhang Q, Han X-Z, Burraco P, Hao X, Teng L-W, Liu Z-S, et al. Telomere length, oxidative stress and their links with growth and survival in a lizard facing climate warming. Sci Total Environ. 2023;891: 164424. doi:10.1016/j.scitotenv.2023.164424

30. Eastwood JR, Connallon T, Delhey K, Hall ML, Teunissen N, Kingma SA, et al. Hot and dry conditions predict shorter nestling telomeres in an endangered songbird: Implications for population persistence. Proc National Acad Sci. 2022;119: e2122944119. doi:10.1073/pnas.2122944119

31. Ton R, Boner W, Raveh S, Monaghan P, Griffith SC. Effects of heat waves on telomere dynamics and parental brooding effort in nestlings of the zebra finch (Taeniopygia castanotis) transitioning from ectothermy to endothermy. Mol Ecol. 2023. doi:10.1111/mec.17064

32. Cunningham SJ, Gardner JL, Martin RO. Opportunity costs and the response of birds and mammals to climate warming. Front Ecol Environ. 2021;19: 300–307. doi:10.1002/fee.2324

33. Mertens JAL. Thermal conditions for successful breeding in Great Tits (Parus major L.). Oecologia. 1977;28: 1–29. doi:10.1007/bf00346834

34. Ruuskanen S, Espín S, Sánchez-Virosta P, Sarraude T, Hsu B-Y, Pajunen P, et al. Transgenerational endocrine disruption: Does elemental pollution affect egg or nestling thyroid hormone levels in a wild songbird? Environ Pollut. 2019;247: 725–735. doi:10.1016/j.envpol.2019.01.088

35. Espin, Sanchez-Virosta, AJ G-F, T E. A microplate adaptation of the thiobarbituric acid reactive substances assay to determine lipid peroxidation fluorometrically in sm all sample volumes. Revista de Toxicología. 2017;34: 94–98.

36. Stier A, Hsu B-Y, Marciau C, Doligez B, Gustafsson L, Bize P, et al. Born to be young? Prenatal thyroid hormones increase early-life telomere length in wild collared flycatchers. Biology Letters. 2020;16: 20200364–4. doi:10.1098/rsbl.2020.0364

37. Ellegren H, Fridolfsson AK. Male-driven evolution of DNA sequences in birds. Nature genetics. 1997;17: 182–184. doi:10.1038/ng1097-182

38. Chang H-W, Cheng C-A, Gu D-L, Chang C-C, Su S-H, Wen C-H, et al. High-throughput avian molecular sexing by SYBR green-based real-time PCR combined with melting curve analysis. BMC Biotechnology. 2008;8: 12–8. doi:10.1186/1472-6750-8-12

39. Hasselblad V, Hedges LV. Meta-Analysis of Screening and Diagnostic Tests. Psychol Bull. 1995;117: 167–178. doi:10.1037/0033-2909.117.1.167

40. Sauve D, Friesen VL, Charmantier A. The Effects of Weather on Avian Growth and Implications for Adaptation to Climate Change. Frontiers in Ecology and Evolution. 2021;9: 5. doi:10.3389/fevo.2021.569741

41. Bano A, Dhana K, Chaker L, Kavousi M, Ikram MA, Mattace-Raso FUS, et al. Association of Thyroid Function With Life Expectancy With and Without Cardiovascular Disease. JAMA Internal Medicine. 2017;177: 1650–8. doi:10.1001/jamainternmed.2017.4836

42. Rey B, Romestaing C, Bodennec J, Dumet A, Fongy A, Duchamp C, et al. Thyroid status affects membranes susceptibility to free radicals and oxidative balance in skeletal muscle of Muscovy ducklings (Cairina moschata). Journal of Experimental Zoology Part A: Ecological Genetics and Physiology. 2014; n/a-n/a. doi:10.1002/jez.1872

43. Xie S, Tearle R, McWhorter TJ. Heat Shock Protein Expression is Upregulated after Acute Heat Exposure in Three Species of Australian Desert Birds. Avian Biology Research. 2019;11: 263–273. doi:10.3184/175815618×15366607700458

44. Lattin CR, Romero LM. Chronic stress alters concentrations of corticosterone receptors in a tissue-specific manner in wild house sparrows (Passer domesticus). J Exp Biol. 2014;217: 2601–2608. doi:10.1242/jeb.103788

45. Pradhan DS, Ness RV, Jalabert C, Hamden JE, Austin SH, Soma KK, et al. Phenotypic flexibility of glucocorticoid signaling in skeletal muscles of a songbird preparing to migrate. Horm Behav. 2019;116: 104586. doi:10.1016/j.yhbeh.2019.104586

46. Hafen PS, Abbott K, Bowden J, Lopiano R, Hancock CR, Hyldahl RD. Daily heat treatment maintains mitochondrial function and attenuates atrophy in human skeletal muscle subjected to immobilization. Journal of applied physiology (Bethesda, Md : 1985). 2019;127: 47–57. doi:10.1152/japplphysiol.01098.2018

47. Fiorenza M, Lemminger AK, Marker M, Eibye K, Iaia FM, Bangsbo J, et al. High-intensity exercise training enhances mitochondrial oxidative phosphorylation efficiency in a temperature-dependent manner in human skeletal muscle: implications for exercise performance. Faseb J. 2019;33: 8976–8989. doi:10.1096/fj.201900106rrr

48. Stier A, Bize P, Hsu B-Y, Ruuskanen S. Plastic but repeatable: rapid adjustments of mitochondrial function and density during reproduction in a wild bird species. Biology Letters. 2019;15: 20190536. doi:10.1098/rsbl.2019.0536

49. Salmón P, Nilsson JF, Watson H, Bensch S, Isaksson C. Selective disappearance of great tits with short telomeres in urban areas. Proceedings of the Royal Society of London Series B: Biological Sciences. 2017;284: 20171349–8. doi:10.1098/rspb.2017.1349

50. Martens DS, Plusquin M, Cox B, Nawrot TS. Early Biological Aging and Fetal Exposure to High and Low Ambient Temperature: A Birth Cohort Study. Environmental Health Perspectives. 2019;127: 117001–10. doi:10.1289/ehp5153

51. Ni W, Wolf K, Breitner S, Zhang S, Nikolaou N, Ward-Caviness CK, et al. Higher Daily Air Temperature Is Associated with Shorter Leukocyte Telomere Length: KORA F3 and KORA F4. Environ Sci Technol. 2022;56: 17815–17824. doi:10.1021/acs.est.2c04486

52. Casagrande S, Stier A, Monaghan P, Loveland JL, Boner W, Lupi S, et al. Increased glucocorticoid concentrations in early life cause mitochondrial inefficiency and short telomeres. Journal Of Experimental Biology. 2020;223: jeb222513–14. doi:10.1242/jeb.222513

53. Dupoué A, Marciau C, Miles D, Meylan S. Shorter telomeres precede population extinction in wild lizards. Scientific Reports. 2017;7: 1–9. doi:10.1038/s41598-017-17323-z

## References

1. Aljanabi SM, Martinez I. Universal and rapid salt-extraction of high quality genomic DNA for PCR-based techniques. Nucleic Acids Res. 1997;25: 4692–4693. doi:10.1093/nar/25.22.4692

2. Larsen S, Nielsen J, Hansen CN, Nielsen LB, Wibrand F, Stride N, et al. Biomarkers of mitochondrial content in skeletal muscle of healthy young human subjects. The Journal of Physiology. 2012;590: 3349–3360. doi:10.1113/jphysiol.2012.230185

3. Stier A, Bize P, Hsu B-Y, Ruuskanen S. Plastic but repeatable: rapid adjustments of mitochondrial function and density during reproduction in a wild bird species. Biology Letters. 2019;15: 20190536. doi:10.1098/rsbl.2019.0536

4. Atema E, Mulder E, Noordwijk AJV, Verhulst S. Ultralong telomeres shorten with age in nestling great tits but are static in adults and mask attrition of short telomeres. Molecular Ecology Resources. 2019;19: 648–658. doi:10.1111/1755-0998.12996

5. Stier A, Massemin S, Zahn S, Tissier ML, Criscuolo F. Starting with a handicap: effects of asynchronous hatching on growth rate, oxidative stress and telomere dynamics in free-living great tits. Oecologia. 2015;179: 999–1010. doi:10.1007/s00442-015-3429-9

6. Stier A, Delestrade A, Bize P, Zahn S, Criscuolo F, Massemin S. Investigating how telomere dynamics, growth and life history covary along an elevation gradient in two passerine species. Journal of Avian Biology. 2016;47: 134–140. doi:10.1111/jav.00714

7. Salmon P, Nilsson JF, Nord A, Isaksson C. Urban environment shortens telomere length in nestling great tits, Parus major. Biology Letters. 2016; 1–4. doi:10.1098/rsbl.2016.0155&domain=pdf&date_stamp=2016-06-14

8. Salmón P, Nilsson JF, Watson H, Bensch S, Isaksson C. Selective disappearance of great tits with short telomeres in urban areas. Proceedings of the Royal Society of London Series B: Biological Sciences. 2017;284: 20171349–8. doi:10.1098/rspb.2017.1349

9. Kärkkäinen T, Briga M, Laaksonen T, Stier A. Within-individual repeatability in telomere length: A meta-analysis in nonmammalian vertebrates. Mol Ecol. 2021. doi:10.1111/mec.16155

10. Ellegren H, Fridolfsson AK. Male-driven evolution of DNA sequences in birds. Nature genetics. 1997;17: 182–184. doi:10.1038/ng1097-182

11. Chang H-W, Cheng C-A, Gu D-L, Chang C-C, Su S-H, Wen C-H, et al. High-throughput avian molecular sexing by SYBR green-based real-time PCR combined with melting curve analysis. BMC Biotechnology. 2008;8: 12–8. doi:10.1186/1472-6750-8-12

12. Casagrande S, Stier A, Monaghan P, Loveland JL, Boner W, Lupi S, et al. Increased glucocorticoid concentrations in early life cause mitochondrial inefficiency and short telomeres. Journal Of Experimental Biology. 2020;223: jeb222513–14. doi:10.1242/jeb.222513

13. Verhagen I, Laine VN, Mateman AC, Pijl A, Wit R de, Lith B van, et al. Fine-tuning of seasonal timing of breeding is regulated downstream in the underlying neuro-endocrine system in a small songbird. Journal Of Experimental Biology. 2019;222: jeb202481–22. doi:10.1242/jeb.202481

14. Criscuolo F, Bize P, Nasir L, Metcalfe NB, Foote CG, Griffiths K, et al. Real-time quantitative PCR assay for measurement of avian telomeres. J Avian Biol. 2009;40: 342–347. doi:10.1111/j.1600-048x.2008.04623.x

15. Bates D, Mächler M, Bolker B, Walker S. Fitting Linear Mixed-Effects Models Using lme4. J Stat Softw. 2015;67. doi:10.18637/jss.v067.i01

16. Halekoh U, Højsgaard S. A Kenward-Roger Approximation and Parametric Bootstrap Methods for Tests in Linear Mixed Models - The R Package pbkrtest. J Stat Softw. 2014;59. doi:10.18637/jss.v059.i09

